# Lack of neuroprotection after systemic administration of the soluble TNF inhibitor XPro1595 in an rAAV6-α-Syn+PFFs-induced rat model for Parkinson’s disease

**DOI:** 10.1101/2024.10.23.619575

**Authors:** Filip Fredlund, Claes Fryklund, Olivia Trujeque-Ramos, Hannah A. Staley, Joaquin Pardo, Kelvin C. Luk, Malú G. Tansey, Maria Swanberg

**Affiliations:** Translational Neurogenetics Unit, Department of Experimental Medical Science, Lund University, Lund, Sweden; Department of Neuroscience, University of Florida College of Medicine, Gainesville, FL, USA; Instituto de Investigaciones Bioquímicas de La Plata “Profesor Doctor Rodolfo R. Brenner”, Facultad de Ciencias Médicas, Universidad Nacional de La Plata, Buenos Aires, Argentina; Molecular Neuromodulation, Wallenberg Neuroscience Center, Lund University, Lund, Sweden; Department of Pathology and Laboratory Medicine, Center for Neurodegenerative Disease Research, University of Pennsylvania Perelman School of Medicine, Philadelphia, PA, USA

**Keywords:** TNF, Parkinson’s disease, neurodegeneration, neuroinflammation, α-Synuclein, PFF, CIITA

## Abstract

Parkinson’s disease (PD) is characterized by dopaminergic neurodegeneration, α-Synuclein (α-Syn) pathology, and inflammation. Microglia in the substantia nigra pars compacta (SNpc) upregulate major histocompatibility complex class II (MHCII), and variants in genes encoding MHCII affect PD risk. Additionally, elevated TNF levels and α-Syn-reactive T cells in circulation suggest a strong link between innate and adaptive immune responses in PD. We have previously reported that reduced levels of the class II transactivator, the master regulator of MHCII expression, increases susceptibility to α-Syn-induced PD-like pathology in rats and are associated with higher serum levels of soluble TNF (sTNF). Here, we demonstrate that inhibiting sTNF with a dominant-negative TNF variant, XPro1595, known to be neuroprotective in endotoxin- and toxin-induced neurodegeneration models, fails to protect against robust α-Syn-induced PD-like pathology in rats. We used a model combining rAAV-mediated α-Syn overexpression in SNpc with striatal injection of α-Syn preformed fibrils two weeks later. Systemic XPro1595 treatment was initiated one-week post-rAAV-α-Syn. We observed up to 70% loss of striatal dopaminergic fibers without treatment, and no protective effects on dopaminergic neurodegeneration after XPro1595 administration. Pathological α-Syn levels as well as microglial and astrocytic activation were not reduced in SNpc or striatum following XPro1595 treatment. An increase in IL-6 and IL-1β levels in CSF was observed in rats treated with XPro1595, possibly explaining a lack of protective effects following treatment. Our results highlight the need to determine the importance of timing of treatment initiation, which is crucial for future applications of sTNF therapies in PD patients.

## Introduction

Idiopathic Parkinson’s disease (PD) is a multifactorial neurodegenerative disease, characterized by progressive loss of dopaminergic neurons in the substantia nigra pars compacta (SNpc), intraneuronal inclusions of α-Synuclein (α-Syn) and neuroinflammation (Poewe et al., 2017). Currently, there are no disease-modifying medications available for PD, causing affected individuals to experience progressive symptoms and disability (Poewe et al., 2017).

The inflammatory components in PD are complex and studies have highlighted the importance of both systemic inflammation and neuroinflammation on disease susceptibility and progression (Harms et al., 2021; Harms et al., 2023; Hirsch and Hunot, 2009; Tansey et al., 2022). The human leukocyte antigen (*HLA*) region encodes major histocompatibility complex class II (MHCII) proteins that present antigens to CD4+ T cells, initiating an adaptive immune response. Genetic association between PD and single nucleotide variants in the *HLA* region points out antigen presentation and adaptive immunity as potential contributors to PD (Hamza et al., 2010; Kannarkat et al., 2015; Wissemann et al., 2013; Yu et al., 2021). In support of this, studies have found α-Syn reactive CD4+ T cells in the circulation of individuals with PD (Gate et al., 2021; Lindestam Arlehamn et al., 2020; Sulzer et al., 2017), and several proinflammatory cytokine levels have been found to be altered in PD, including soluble tumor necrosis factor (sTNF) (Qu et al., 2023). Targeting the inflammatory response in PD has been proposed to be a viable treatment strategy to mitigate PD susceptibility and progression (Grotemeyer et al., 2022; Hirsch and Hunot, 2009; Weiss et al., 2022).

We and others have proposed the major regulator of MHCII expression, the class II transactivator (CIITA), as a target of interest to alter PD progression (Fredlund et al., 2024; Jimenez-Ferrer et al., 2021; Jimenez-Ferrer et al., 2017; Williams et al., 2018). In a recombinant adeno associated viral vector (rAAV) -α-Syn-induced *in vivo* mouse model of PD, both global knock-out and local silencing of the *Ciita* gene were reported to prevent neurodegeneration, MHCII expression and infiltration of peripheral immune cells (Williams et al., 2018). In contrast, we have found in rAAV-α-Syn- and rAAV-α-Syn+α-Syn preformed fibrils (PFFs) -induced PD models that congenic rats (DA.VRA4) with naturally occurring reduced *Ciita* expression had increased susceptibility to neurodegeneration, pathological α-Syn spread and motor dysfunction. We also found that lower *Ciita* and MHCII expression levels were associated with a more inflammatory-prone state, likely related to an increased number of MHCII+ microglia (Fredlund et al., 2024; Jimenez-Ferrer et al., 2021; Jimenez-Ferrer et al., 2017). Of great interest, congenic DA.VRA4 rats have constitutively higher levels of sTNF in serum, which might impact the susceptibility to α-Syn-induced PD-like pathology (Fredlund et al., 2024).

TNF inhibitors are used in a clinical setting today to treat immune-related disorders, including rheumatoid arthritis and Crohn’s disease. These inhibitors target both sTNF and transmembrane TNF (tmTNF). sTNF signals through TNF receptor 1 (TNFR1) found on most cells whereas tmTNF signals through both TNFR1 and TNF receptor 2 (TNFR2), which is found on immune cells, epithelial cells and cells in the CNS (Probert, 2015). Although both sTNF/TNFR1 and tmTNF/TNFR2 signaling induce a proinflammatory response, tmTNF/TNFR2 signaling in the CNS is pro-survival, neuroprotective, and important for remyelination of demyelinated neurons (Probert, 2015). Therefore, selective inhibition of sTNF has been proposed for treating PD. Such sTNF inhibition using dominant-negative TNF (DN-TNF) (Steed et al., 2003) has successfully protected against dopaminergic neurodegeneration and mitigating the neuroinflammatory response in toxin-induced models (Barnum et al., 2014; Harms et al., 2011; McCoy et al., 2006; McCoy et al., 2008). However, the approach has, to our knowledge, not previously been tested in models with α-Syn-induced PD-like pathology.

Here, we aim to investigate if systemic administration of DN-TNF, using XPro1595, can protect against α-Syn-induced PD-like neurodegeneration and neuroinflammation. As a model system, we used the rAAV-α-Syn+PFFs induced PD model in the DA.VRA4 rat strain that displays an inflammatory-prone steady-state, partly characterized by elevated serum sTNF levels. The unilateral combined rAAV-α-Syn+PFFs-induced PD model shares several key features with PD including α-Syn-induced progressive dopaminergic neurodegeneration, α-Syn pathology, and an inflammatory response, which can be initiated at a lower virus titer compared to using AAV-mediated α-Syn overexpression alone (Bjorklund et al., 2022; Cenci and Bjorklund, 2020). Our results show that systemic treatment with the sTNF inhibitor XPro1595 reaches the CNS, but fails to exert neuroprotective effects, limit α-Syn pathology, or reduce the inflammatory response in the rAAV-α-Syn+PFFs model.

## Materials and methods

### Study design

A total of 20 male DA.VRA4 rats, at 12 ± 3 weeks of age, were unilaterally injected with 3 µl rAAV6-α-Syn (1.3E+10 gc/µl) in the SNpc at the start (week 0) of the study (Fig. 1A). Starting one week later, rats received s.c. injections in the flank, with saline (n = 10) or the sTNF inhibitor XPro1595 (10 mg/kg) (n = 10), every third day for the duration of the study (a total of 17 injections/rat). At week 2, all rats were unilaterally (same hemisphere as for rAAV6-α-Syn injections) injected with 3 µl sonicated PFFs (2.5 µg/µl) in the dorsal striatum. All rats were euthanized eight weeks after SNpc injection and samples were collected for future analyses. All analyses were done in a blinded manner. Two rats from the saline treated group were excluded due to missed SN during rAAV6-α-Syn injection. Tissue from two DA.VRA4 rats injected with 3 µl rAAV6-(-) (human α-Syn transgene excised, 1.7E+10 gc/µl) and 3 µl of DBPS two weeks later as previously described (Fredlund et al., 2024) were included as indicative controls for the α-Syn-induced PD model (Supplementary Fig. 1).

**Figure 1.**
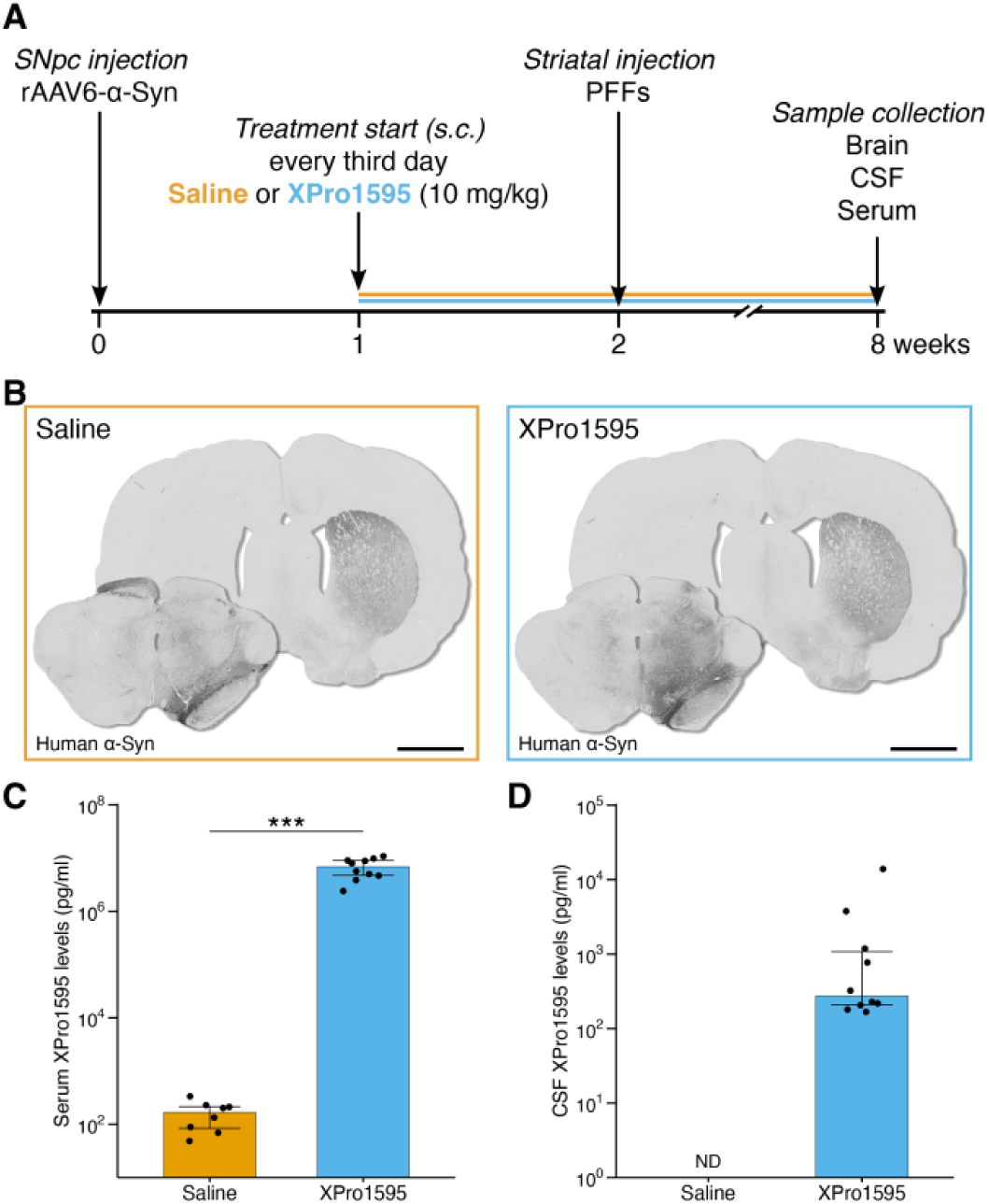
Experimental timeline, confirmation of human α-Syn expression in SNpc and striatum, and XPro1595 measurments in serum and CSF. A. Experimental outline. B. Human α-Syn expression eight weeks after unilateral rAAV-α-Syn injection in SNpc combined with striatal seeding with PFFs. Scale bar = 2 mm. C. Bar graph showing XPro1595 (and background signal) concentration on a log_10_ scale in serum. Median and IQR with individual values are shown. Statistical analysis was conducted using Mann-Whitney U test. ***p < 0.001. D. Bar graph showing XPro1595 levels on a log_10_ scale in CSF. Median and IQR with individual values. Group numbers; Saline n = 8, XPro1595 n = 10. Abbreviations. SNpc, substantia nigra pars compacta; rAAV6, recombinant adeno-associated virus serotype 6; α-Syn, α-Synuclein; s.c., subcutaneous; PFFs, preformed fibrils; CSF, cerebrospinal fluid; ND, non-detected; sTNF, soluble tumor necrosis factor; IQR, interquartile range.

### Animals

DA.VRA4 founders were donated by Professor Fredrik Piehl. The congenic DA.VRA4 strain was generated by transfer of the VRA4 locus from the PVG strain to a DA strain background following at least 10 generations of backcrossing (Harnesk et al., 2008). 2-3 rats were housed in individually ventilated cages (type III high) with water and standard rodent chow *ad libitum*, and kept in a pathogen-free and climate-controlled environment with a 12 h light/dark cycle at the Biomedical Center in Lund. All experimental animal procedures performed in this project was carried out in accordance with current EU legislation and were approved by the local ethics committee in the Malmö - Lund region and specified in permit 18037/2019 and in the amendment 15349/2022.

### Viral vector

rAAV6 vector carrying human α-Syn under transcriptional regulation by the Synapsin-1 promotor and the woodchuck hepatitis virus posttranscriptional regulatory element (Decressac et al., 2012), referred to as rAAV6-α-Syn. The rAAV6-α-Syn vector with the human α-Syn excised was used as control, referred to as rAAV6-(-) (Fredlund et al., 2024). The rAAV6 vectors were produced by the AAV platform, formerly part of the strategic research area MultiPark, as previously described (Decressac et al., 2011). The concentration of the rAAV6 vectors was determined by ITR-qPCR.

### Preformed fibrils

Human α-Syn PFFs were produced as previously described (Luk et al., 2009) and stored at -80°C until use. PFFs were diluted to a concentration of 2.5 µg/µl using sterile Dulbecco’s phosphate buffered saline (DPBS) and sonicated for 6 min with 1 s on/1 s off pulses at 70% power using a Q125 sonicator fitted with a cup horn (Qsonica, USA) immediately before use. DPBS was used as vehicle control in the control group.

### Stereotaxic surgical procedure

Stereotaxic surgery for rAAV6-α-Syn/rAAV6-(-) SNpc and striatal PFFs/DPBS injection was done as previously described (Fredlund et al., 2024), with a minor adjustment where sutures were used instead of surgical staples. Injections were made into the right hemisphere (ipsilateral). The following coronal coordinates relative to Bregma was used for rAAV6-α-Syn SNpc injection; rostral/caudal (R/C) -5.3 mm, medial/lateral (M/L) ± 1.7 mm and dorsal/ventral (D/V) -7.2 mm (Paxinos and Watson, 2014). For injections of PFFs into the ipsilateral dorsal striatum, the following coordinates, relative to Bregma, were used; R/C -0.4 mm, M/L ± 0.3 mm, and D/V -4.5 mm (Paxinos and Watson, 2014).

### Tissue collection

Rats were euthanized by intraperitoneal injection of 200-300 mg/kg sodium pentobarbital (APL, Sweden). CSF, serum, and brain collection (PFA perfused) was done as previously described (Fredlund et al., 2024). Here follows a brief description of each tissue collection.

CSF samples were collected using a 27G small scalp vein set (Vygon, France) from an exposed cisterna magna from rats attached to a stereotaxic frame with an approximate 50-60° downward flex of the head. Samples were collected into protein LoBind tubes (Eppendorf, Germany) and stored at -80°C until analysis.

Whole blood collected from cardiac puncture was left undisturbed at RT for 45-60 min followed by 10 min centrifugation at 4°C and 2,000xg. The separated serum was aliquoted into protein LoBind tubes (Eppendorf. Germany) and stored at -80°C until analysis.

Rats were transcardially perfused, with the descending aorta clamped, with 0.9% (w/v) saline for at least 5 min, followed by 5 min perfusion with ice-cold 4% PFA (w/v). Brains were post-fixed in 4% PFA (w/v) at 4°C overnight followed by cryopreservation in PBS (pH 7.2) with 30% sucrose (w/v) and 0.01% sodium azide (w/v) until sectioning.

### XPro1595 and cytokine measurements

XPro1595 levels in serum and CSF were measured using the “V-PLEX Human TNF-α Kit” (K151QWD) with an adapted protocol; the calibrator included in the kit was substituted with XPro1595. A standard curve was generated by six 1:4 serial dilutions of XPro1595, from 250,000 pg/ml to 61 pg/ml. A blank was also included in the standard curve. Other than modifying the calibrator, the plate was run according to manufacturers’ instructions (MSD, USA). Serum samples were diluted 1:200 and CSF samples were run undiluted. Calibrator (XPro1595) and samples were all run in duplicates.

Multiplexed ELISA of nine soluble cytokines (IFN-γ, IL-1β, IL-4, IL-5, IL-6, IL-10, CXCL1, and TNF) was measured on neat serum and CSF samples using the “V-PLEX Proinflammatory Panel 2 Rat Kit” (K15059) according to the manufacturers’ instructions (MSD, USA). Calibrator and serum samples were run in duplicates. Due to limited sample volume remaining after XPro1595 measurements, CSF samples were run in singlets. TNF measurements were excluded from analysis since XPro1595 may be detected as native TNF. If more than one sample/group for a cytokine and sample type (CSF or serum) was non-detected (ND), results are reported as “ND” at group level. If only one sample was ND for a specific cytokine, sample type, and group, it was replaced by the lowest quantifiable value divided by two.

All MSD plates were read on a MESO QuickPlex SQ 120 analyzer (MSD, USA) and analyzed using the Discovery Workbench software version 4.0.13 (MSD, USA) to convert the signal to pg/ml.

### Immunohistochemistry

Sectioning and 3,3’-diaminobenzidine (DAB) staining on free floating sections was done as previously described (Fredlund et al., 2024). Sections were collected in series of 12. 35 µm coronal sections were quenched with 3% H_2_O_2_ (v/v) (Supelco, USA) and 10% methanol (v/v) (J.T. Baker, USA) in PBS (pH 7.2). Blocking was done using 10% normal serum (Biowest, France) in 0.3% PBST. Primary antibody was diluted in 0.3% PBST containing 5% normal serum (Biowest, France) and incubated at 4°C overnight. Secondary antibody was diluted in 0.3% PBST and incubated for 2 h at RT. Horseradish peroxidase conjugated avidin/biotin-complex (Vector laboratories, USA) and DAB substrate kit (Vector laboratories, USA) were used according to manufacturers’ instructions. A detailed overview of serum and antibodies used for immunohistochemistry (IHC) can be found in table 1. Sections were dehydrated and coverslip attached using Pertex (Histolab, Sweden).

**Table 1.**
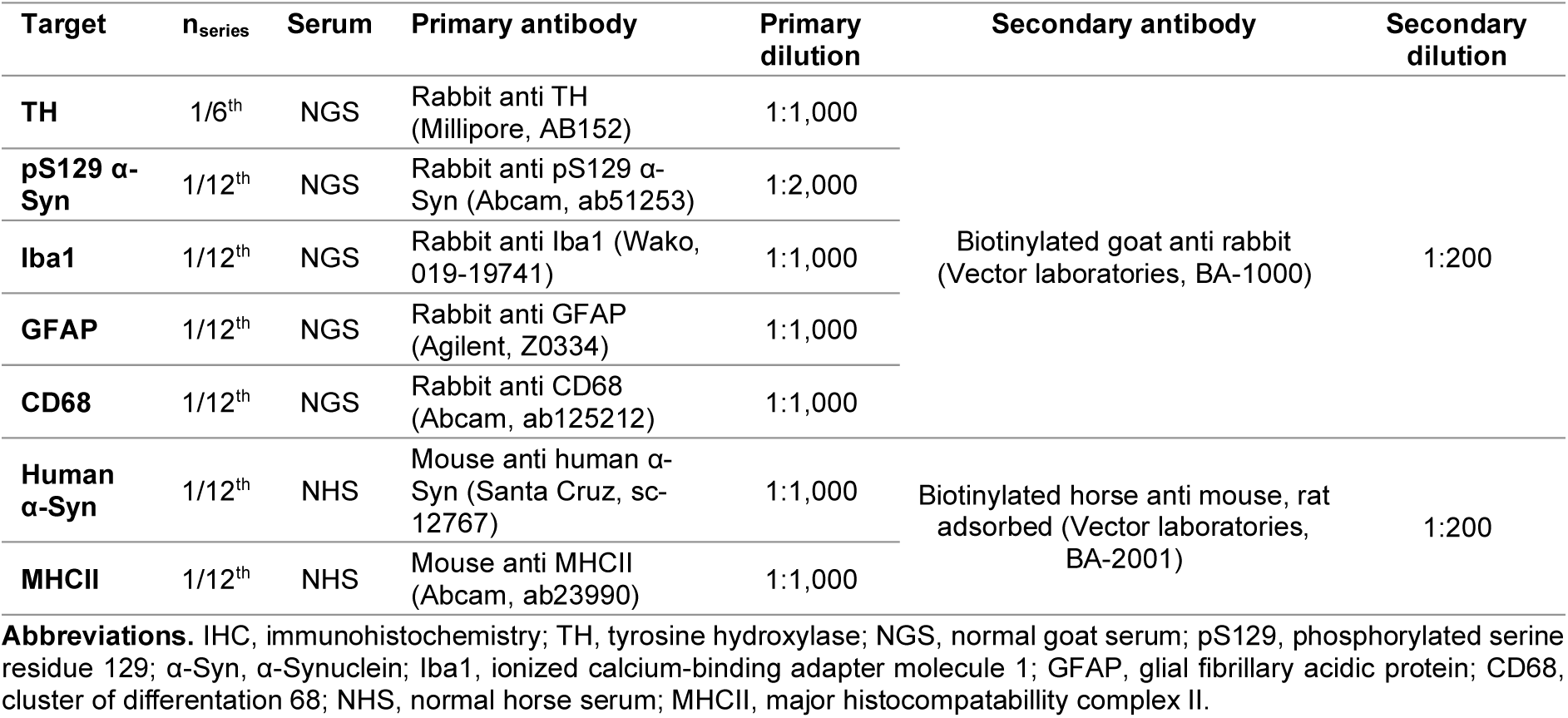
List of number of series, blocking reagents and antibodies with their corresponding dilution used for IHC.

### Microscopy

All images were acquired using an Olympus BX61 with a VS-120 automated stage (Olympus, Japan). For images captured at 2x and 4x magnification a single z-plane image was acquired. For images captured at 10x and 20x, z-planes spaced 1.5 µm apart, were acquired and collapsed to a single focal plane using extended focal imaging using the VS-ASW-S6 software version 2.9 (Olympus, Japan).

### Deep convolutional neural network algorithm models for IHC analyses

AI models using deep convolutional neural network (CNN) algorithms with supervised learning was developed using aiforia create (Aiforia technologies, Finland). Images, acquired at 20x magnification, were uploaded to the aiforia cloud (Aiforia technologies, Finland). A summary of the AI models can be found in table 2.

**Table 2.** AI models used for IHC analysis. Summary of the AI training and performance of the three models developed for IHC analysis. For layers with object & IS, precision, recall and F1 scores refer to performance of object detection (metrics not available for IS).

| Model (iterations) | AI engine version | Layer type | FOV/object size ($\mu\text{m}$) | Complexity | Annotations | Precision (%) | Recall (%) | F1 score (%) |
| --- | --- | --- | --- | --- | --- | --- | --- | --- |
| TH+ SNpc cell (700) | 5 | Object | 20 | Very Complex | 269 objects across 56 regions | 92.31 | 98.14 | 95.14 |
| TH+ axonal swelling (1,000) | 7 | Region (parent) | 1,000 | Very Complex | 114 | 100 | 99.94 | 99.97 |
|  |  | Object & IS (child) | 5 | Intermediate (object) & Complex (IS) | 405 objects across 224 regions | 97.67 | 95.45 | 96.55 |
| Iba1+ cell (2,193) | 7 | Region (parent) | 100 | Very Complex | 41 | 99.49 | 98.14 | 98.81 |
|  |  | Object & IS (child) | 40 | Intermediate (object) & Ultra Complex (IS) | 235 objects across 231 regions | 98.94 | 91.18 | 94.40 |

**Abbreviations.** AI, Artificial intelligence; IHC, immunohistochemistry; IS, instance segmentation; FOV, field of view; TH, tyrosine hydroxylase; SNpc, substantia nigra pars compacta; Iba1, ionized calcium-binding adapter molecule 1.

The TH+ SNpc cell model was trained to identify TH+ cells of diverse morphologies and staining intensities in SNpc.

The TH+ axonal swelling model was developed to identify swellings in the dorsal striatum. The parent layer for the TH+ axonal swelling model was trained to recognize brain tissue of very diverse staining intensities of brain tissue. To enable identification of TH+ axonal swellings of various sizes, instance segmentation was utilized to manually outline every TH+ axonal swelling object (child layer), using a training data set that was not included in formal analyses. TH+ axonal swellings annotated for training were approximately >3 µm in diameter, a size that were almost devoid in the contralateral hemisphere.

The Iba1+ cell model was developed to identify Iba1+ cells in dorsal striatum and SN. The parent layer was trained to identify brain tissue of diverse complexities. To enable more complex analysis at a cellular level, instance segmentation was used to outline entire Iba1+ cell objects (child layer). The AI was trained on a data set that was not included in formal analyses. Post-training parameters for instance segmentation were altered with the following settings; gain factor set to 1.5, level of granularity details was set to high, and “prevent instance overlap” was enabled.

### Quantification of TH+ SNpc cells

Coronal midbrain sections between R/C coordinates (relative to Bregma) -4.56 and -6.6 (Paxinos and Watson, 2014), captured at 20x, was used for analyses. For quantification of TH+ SNpc cells every contra- and ipsilateral SNpc region was manually annotated and analyzed using the developed TH+ SNpc cell AI model. A total of 7-9 SNpc sections were quantified/rat injected with rAAV6-α-Syn+PFFs. A total of 3-6 SNpc sections were quantified/rat injected with rAAV-(-)+DPBS. The mean TH+ cell counts for contra- and ipsilateral SNpc per rat was normalized to the corresponding group mean contralateral TH+ cell count and expressed as percent remaining TH+ cells in SNpc.

### TH+ fiber density

Coronal sections, captured at 10x and matched at the following ten approximate R/C coordinates (relative to Bregma) was used for TH+ fiber density analysis; 1.6, 1.2, 0.84, 0.36, 0.0, -0.4, -0.8, -1.2, -1.6, and -2.04 in the rAAV6-α-Syn+PFFs group (Paxinos and Watson, 2006; Paxinos and Watson, 2014). Five sections, captured at 4x and matched on the following Bregma coordinates were used for quantification in the rAAV-(-)+DPBS group; 1.6, 0.84, 0.0, -0.8, and -1.6 (Paxinos and Watson, 2006; Paxinos and Watson, 2014). Images were analyzed using Fiji with ImageJ2 version 2.14.0. Images were converted to 8-bit and TH+ fiber density was calculated by inverted mean grey values in contra- and ipsilateral dorsal striatum. Background signal measured from cortex was subtracted in analysis. Ipsilateral TH+ fiber density was expressed as percent relative to the contralateral dorsal striatum for each rat.

### TH+ axonal swellings quantification

A total of five coronal sections/rat, acquired at 20x magnification, matched on the following approximate R/C coordinates (relative to Bregma); 1.2, 0.36, -0.4, -1.2, and -2.04 (1.6, 0.84, 0.0, -0.8, and -1.6 for rAAV6-(-)+DPBS) (Paxinos and Watson, 2006; Paxinos and Watson, 2014), were used to quantify TH+ axonal swellings using the developed TH+ axonal swelling AI model. The sum of all analyzed TH+ axonal swellings were used to generate violin plots to visualize probability densities across hemispheres and treatment. The frequency of TH+ axonal swellings of various sizes, normalized to group number, were used to generate histograms. The number of TH+ axonal swellings per animal was first normalized to striatum area analyzed, and then further normalized to the corresponding group and hemisphere TH+ SNpc cell count.

### pS129 α-Syn, MHCII, CD68, and GFAP quantification

Coronal sections, acquired at 20x magnification, was used for analysis. 5 striatal sections/rat matched on the following approximate R/C coordinates, relative to Bregma, were used; 1.2, 0.36, -0.4, -1.2, and -2.04 (1.6, 0.84, 0.0, -0.8, and -1.6 for rAAV6-(-)+DPBS) (Paxinos and Watson, 2006; Paxinos and Watson, 2014). For SN analysis 4-6 (pS129 α-Syn and GFAP) or 3-5 (MHCII and CD68) sections/rat in the range of -4.56 and -6.6 R/C coordinates, relative to Bregma, were used for quantification (Paxinos and Watson, 2014). For pS129 α-Syn only ipsilateral hemisphere was used for analyses (excluded for rAAV6-(-)+DPBS group), whereas both contra- and ipsilateral hemispheres were included for MHCII, CD68, and GFAP analyses. The percentage positive area for each marker, normalized to region of interest area, were acquired at each coronal level using the open software QuPath (version 0.5.1). The mean percentage of immunoreactive positive area/region for each rat were used to compare groups.

### Iba1+ cell analysis

For analysis in dorsal striatum 4-5 sections/rat, acquired at 20x magnification, matched on the following R/C coordinates relative to Bregma; 1.2, 0.36, -0.4, -1.2, and -2.04 (1.6, 0.84, 0.0, -0.8, and -1.6 for rAAV6-(-)+DPBS) (Paxinos and Watson, 2006; Paxinos and Watson, 2014), were used for quantification. For SN analysis 1-6 sections/rat, between R/C coordinates (relative to Bregma) -4.56 to -6.6, were used for quantification (Paxinos and Watson, 2014). To compare groups and hemispheres, the mean cell characteristic of interest/animal from either dorsal striatum or SN was quantified. To compare Iba1+ cell counts, the mean Iba1+ cell count, normalized to region of interest area, was used for analysis. One saline-treated rat was excluded from analysis due to issues during IHC procedure. For one saline-treated rat there were no intact sections at SN coordinates to be included in analysis.

Ultimately, the group number for Iba1+ cell analysis in the saline group was n = 7 (striatum) and n = 6 (SN).

### Statistical analysis

All generated data was handled in RStudio version 2023.06.0 (Posit PBC, USA). Statistical analyses were also conducted in RStudio. No statistical analysis was performed for the rAAV6-(-)+DPBS group (n=2). Quantile-quantile plots of residuals were used to determine the distribution of data. Mann-Whitney U test was used to compare XPro1595 serum levels and CXCL1 CSF levels between saline and XPro1595 treated rats. Two-way ANOVA was used to test the impact of hemisphere and treatment on the percentage of TH+ SNpc cells, Iba1+ cells (area, area:perimeter and count), percent MHCII+ area, percent CD68+ area, and percentage of GFAP+ area, Tukey’s HSD was used as post-hoc test. Chi-squared goodness-of-fit test was used to compare frequency of TH+ axonal swelling sizes between hemispheres for each treatment group. Unpaired Welch’s t-test was used to compare TH+ axonal swellings, percent positive pS129 α-Syn for ipsilateral hemispheres and cytokine levels with normal distribution between treatment groups. PCA, of Iba1+ cells analyzed in dorsal striatum or SN, was performed using the ‘prcomp’ function, following data standardization. Variable loadings were plotted as vectors on the PCA biplot.

## Results

### Systemic XPro1595 treatment did not protect against rAAV6-α-Syn+PFFs-induced neurodegeneration

rAAV6-α-Syn+PFFs injection resulted in robust unilateral human α-Syn expression (Fig. 1B) which was not observed in the rAAV6-(-)+DPBS control (Supplementary Fig. 1A). XPro1595 was detected in both serum and CSF at sufficient levels to sequester monomeric native sTNF in heterotrimer formations (Steed et al., 2003) following systemic treatment (Fig. 1C and D).

To assess the impact of systemic sTNF inhibition on dopaminergic neurodegeneration we used an AI model to quantify TH+ cells in the SNpc, an approach which has been shown to be comparable to conventional stereology (Penttinen et al., 2018). Both saline and XPro1595 treated rats displayed evident dopaminergic neurodegeneration after unilateral rAAV6-α-Syn+PFFs injections (Fig. 2A). Simple main effects analysis identified an effect of rAAV6-α-Syn+PFFs on loss of TH+ cells in the ipsilateral-compared to contralateral SNpc (F_1, 30_ = 26.2, p < 0.001) in both saline- (-32%, 95% CI [-58, -6.0], p = 0.011) and XPro1595- (-35%, 95% CI [-60, -10], p = 0.0028) treated rats (Fig. 2B). There was no simple main effect of XPro1595 (F_1, 30_ = 0.054, p = 0.82) and no significant interaction (F_1, 30_ = 0.054, p = 0.82) between rAAV6-α-Syn+PFFs (hemisphere) and XPro1595 (treatment) on TH+ cells in the ipsilateral-compared to contralateral SNpc. No dopaminergic cell loss was detected in the rAAV-(-)+DPBS control (Supplementary Fig. 1B).

**Figure 2.**
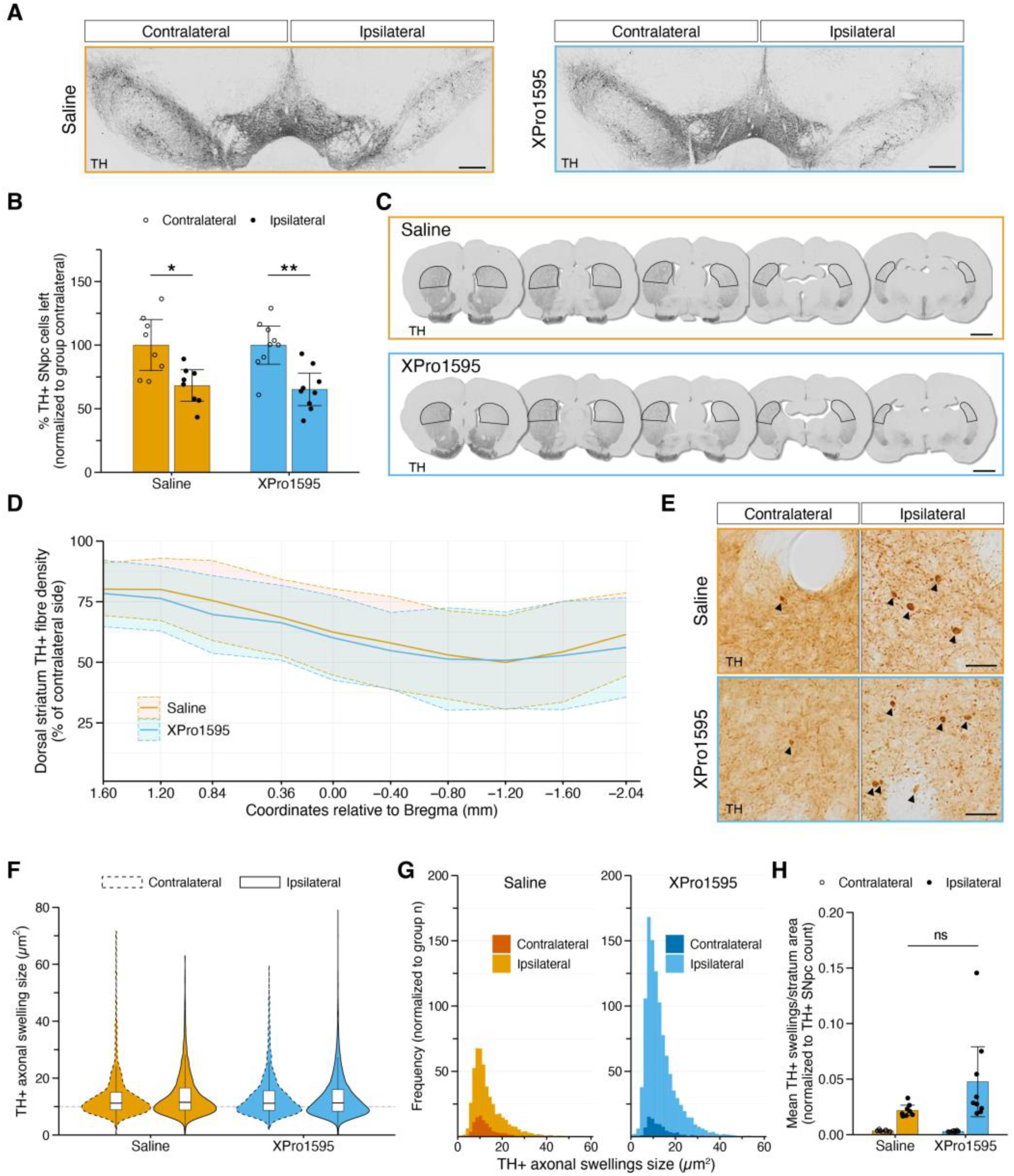
Systemic XPro1595 treatment does not protect against dopaminergic neurodegeneration after combined unilateral rAAV6-α-Syn+PFFs injection. **A.** Representative images of TH+ cells in ventral midbrain of rats treated with saline (left) or XPro1595 (right). Scale bar = 400 µm. **B.** Quantification of the percentage of TH+ cells left in the SNpc normalized to the group contralateral. 7-9 sections/rat was used for quantification. Bar graph showing group mean ± 95% CI with individual values. Statistical analysis was performed using two-way ANOVA, showing an effect of hemisphere (p < 0.001). Post-hoc comparisons were made using Tukey’s HSD test. *p < 0.05, **p < 0.01. **C.** Representative coronal sections stained for TH at various Bregma coordinates in rats treated with saline (top) or XPro1595 (bottom). The TH+ fiber density was measured in the dorsal striatum. Scale bar = 2 mm. **D.** Quantification of the percentage TH+ fiber density remaining in the ipsilateral dorsal striatum after normalization to contralateral hemisphere, plotted against Bregma coordinates along the rostro-caudal axis. 10 sections/rat, matched on Bregma coordinates, was used for analysis. Group mean ± SD is shown. **E.** Representative images of TH+ axonal swellings in the dorsal striatum. Arrowheads indicate axonal swellings identified by the TH+ axonal swelling AI model. Scale bar = 25 µm. **F.** Violin plots of TH+ axonal swelling sizes (µm^2^) based on the sum of TH+ axonal swellings in the dorsal striatum for each hemisphere across groups. **G.** Histogram plots of TH+ axonal swelling sizes (µm^2^) for each hemisphere normalized to group numbers. Statistical analysis was performed to using chi-squared goodness-of-fit test to compare ipsilateral versus contralateral hemispheres (p < 0.001 for both groups) **H.** Quantification of the mean number of TH+ axonal swellings/striatum area normalized to group mean TH+ SNpc count. Group mean ± 95% CI with individual values. Unpaired Welch’s t-test was used to compare treatments in ipsilateral hemispheres (p > 0.05). **F-H.** A total of 5 sections/rat, matched on Bregma coordinates, were used for quantification. Group numbers; saline n = 8 and XPro1595 n= 9. **Abbreviations.** TH, tyrosine hydroxylase; SNpc, substantia nigra pars compacta; ns, non-significant; rAAV6, recombinant adeno-associated virus serotype 6; α-Syn, α-Synuclein; PFFs, preformed fibrils; CI, confidence interval; ANOVA, analysis of variance; HSD, honestly significant difference; SD, standard deviation; AI, artificial intelligence.

In line with TH+ cell loss in SNpc, rAAV6-α-Syn+PFFs resulted in reduced TH+ fiber density in the ipsilateral dorsal striatum of both saline- and XPro1595 treated rats (Fig. 2C). The TH+ fiber density in ipsilateral vs contralateral coronal sections identified similar patterns of axonal neurodegeneration in saline- and XPro1595 treated rats along the rostro-caudal axis (Fig. 2D). No signs of TH+ fiber loss in dorsal striatum was observed in the rAAV6-(-)+DPBS control (Supplementary Fig. 1C).

To investigate if the XPro1595 had an effect on early neurodegenerative processes, we trained an AI model to quantify TH+ axonal swellings in the dorsal striatum (Fig. 2E). The size of TH+ axonal swellings did not differ between ipsilateral and contralateral hemispheres, with the density centering around ∼10 µm^2^ (Fig. 2F and Supplementary Fig. 1D). However, rAAV6-α-Syn+PFFs injections affected the frequency, with significantly more TH+ axonal swellings in the ipsilateral compared to contralateral hemisphere in both saline- (χ^2^49 = Inf, p < 0.001) and XPro1595 (χ^2^49 = Inf, p < 0.001) treated rats (Fig. 2G). No difference in TH+ axonal swelling frequency was observed between ipsi- and contralateral hemispheres in the rAAV6-(-)+DPBS control (Supplementary Fig. 1E). To account for dopaminergic neurodegeneration, TH+ axonal swelling counts were normalized to the number of remaining TH+ cells in the SNpc. There was no difference in normalized TH+ axonal swellings in the ipsilateral hemisphere between saline- and XPro1595 treated groups (0.026, 95% CI [-0.057, 0.0058], p = 0.097) (Fig. 2H).

### Pathologically associated pS129-α-Syn load in the dorsal striatum and in SN was unaffected by systemic XPro1595 administration

pS129 α-Syn is associated with synucleinopathies, including PD (Fujiwara et al., 2002). We investigated if sTNF inhibition by XPro1595 would impact the pathological load of pS129 α-Syn in dorsal striatum and SN.

In both saline- and XPro1595 treated rats, the striatum ipsilateral to rAAV6-α-Syn+PFFs injections had distinct pS129 α-Syn positive signals, mainly observed with a neuritic morphology but also as puncta (Fig. 3A_ii_ and A_iv_). Since there was no apparent signal for pS129 α-Syn in the contralateral hemispheres (Fig. 3A_i_ and A_iii_), only the ipsilateral hemisphere was included for further analyses. There was no difference between saline- and XPro1595 treated rats in pS129 α-Syn positive area normalized to the dorsal striatum area (Fig. 3B).

**Figure 3.**
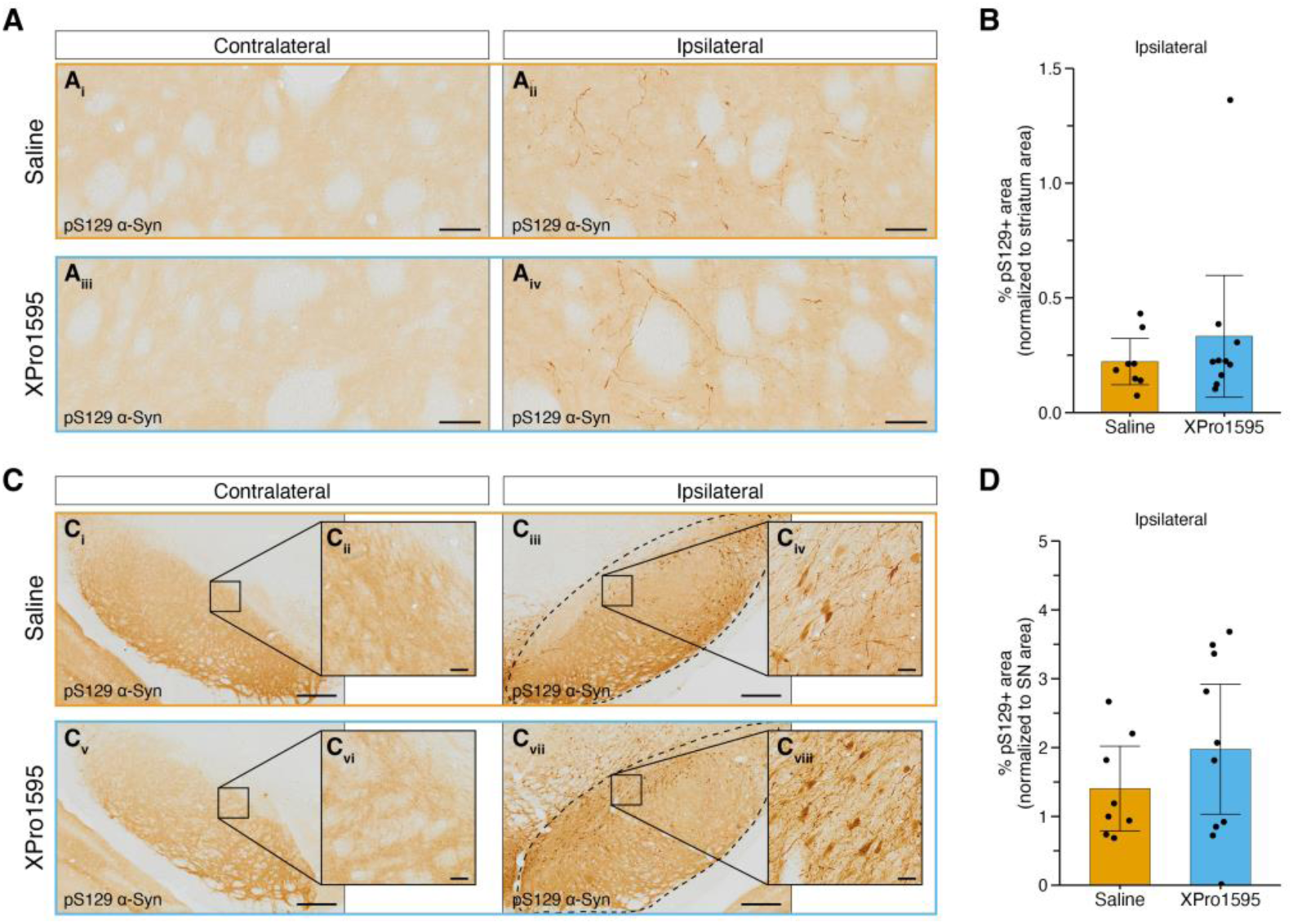
The pathological pS129 α-Syn load is unaffected by systemic XPro1595 treatment. **A.** Representative images of pS129 α-Syn signal in the dorsal striatum in the contralateral and ipsilateral hemispheres in both groups. Scale bar = 50 µm. **B.** Bar graph showing the mean percentage of positive pS129 α-Syn signal normalized to dorsal striatum area, in ipsilateral hemispheres. 5 sections/rat, matched on Bregma coordinates, were used for quantification. Group mean ± 95% CI with individual values. Unpaired Welch’s t-test was used to compare groups. **C.** Representative images of pS129 α-Syn signal in the contralateral and ipsilateral SN in both groups. Representative SN area analyzed is indicated by a dashed line in **C_iii_** and **C_vii_**. Scale bar in **C_i_**, **C_iii_**, **C_v_**, and **C_vii_** = 300 µm. Scale bar in **C_ii_**, **C_iv_**, **C_vi_**, and **C_viii_** = 25 µm. **D.** Quantification of mean percentage positive pS129 α-Syn signal normalized to SN area, in ipsilateral hemispheres. 4-6 sections/rat was used to calculate individual means. Bar graph showing group mean ± 95% CI with individual values. Unpaired Welch’s t-test was used to compare groups. Group numbers; Saline n = 8 and XPro1595 n = 10. **Abbreviations.** pS129, phosphorylated serine residue 129; α-Syn, α-Synuclein; SN, substantia nigra; CI, confidence interval.

Similar to the dorsal striatum, there was no apparent pS129 α-Syn pathology in the contralateral SN (Fig. 3C_i__-ii_ and C_v-vi_). In the SN ipsilateral to rAAV6-α-Syn+PFFs injections, pS129 α-Syn signal was observed within cytoplasm, in neurites and as puncta (Fig. 3C_iii__-iv_ and C_vii-viii_). There was no difference between saline- and XPro1595 treated rats in pS129 α-Syn positive area normalized to SN area (Fig. 3D). No pS129 α-Syn pathology was observed in striatum or SN following rAAV6-(-)+DPBS injection (Supplementary Fig. 1F).

### Neutralization of sTNF with XPro1595 did not mitigate microglial activation in the dorsal striatum or in SN following unilateral rAAV6-α-Syn+PFFs injection

Similar to PD (Harms et al., 2023), the combined rAAV6-α-Syn+PFFs PD model includes activation of microglia, with morphological changes and upregulation of inflammatory markers such as MHCHII and CD68 (Jimenez-Ferrer et al., 2021; Negrini et al., 2022; Thakur et al., 2017). To increase power and minimize user-bias we trained an AI model to detect instance segmentation of Iba1+ cells, enabling both object detection for cell counts and semantic segmentation for cell morphology characteristics. The model efficiently identified Iba1+ cells with various complexity and sizes in dorsal striatum contralateral and ipsilateral to rAAV6-α-Syn+PFFs (Fig. 4A).

**Figure 4.**
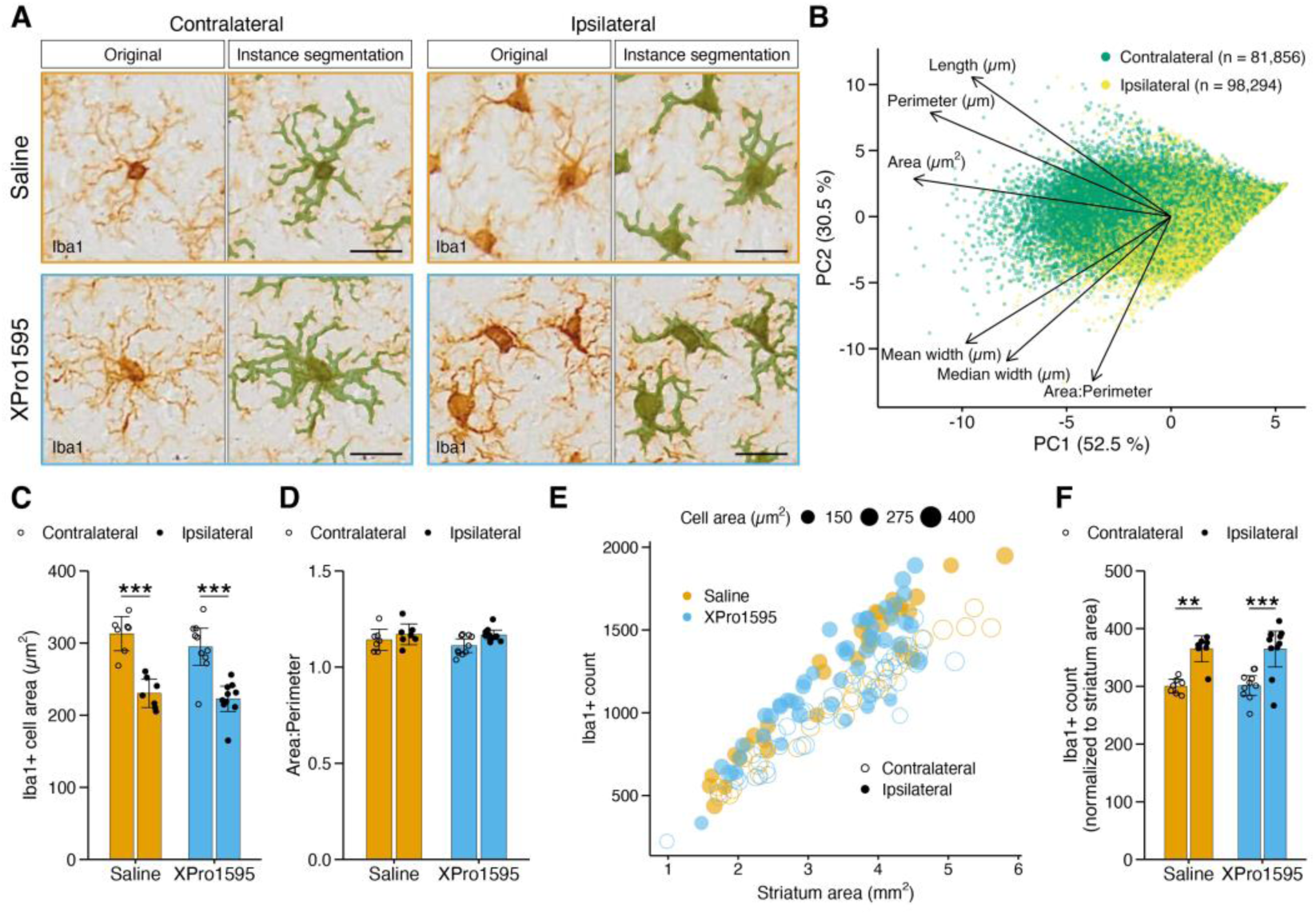
Microglia morphology in dorsal striatum was unaffected by systemic XPro1595 treatmend following combined unilateral injection of rAAV6-α-Syn+PFFs. **A.** Representative images of Iba1+ cells in the contralateral and ipsilateral dorsal striatum in both groups. Instance segmentation overlaps shows how the AI model detected and outlined Iba1+ cells. Scale bar = 20 µm. **B.** PCA biplot with loadings for every instance segmentation event detected in the dorsal striatum, seperated by hemispheres. **C.** Bar graph showing the mean Iba1+ cell area across groups. 4-5 sections/rat, matched on Bregma coordinates, were used to calculate individual means. Group mean ± 95% CI with individual values are shown. Statistical analysis was performed using two-way ANOVA, showing an effect of hemisphere (p < 0.001). Post-hoc comparisons were made using Tukey’s HSD test. ***p < 0.001. **D.** Quantification of the mean ratio of Iba1+ area divided by the Iba1+ perimeter across groups. 4-5 sections/rat, matched on Bregma coordinates, were used to calculate individual means. Bar graph shows mean ± 95% CI with individual values. Two-way ANOVA showed a simple main effect of hemisphere (p < 0.05). Tukey’s HSD was used for post-hoc comparison. **E.** Scatter plot showing the mean Iba1+ cell count against the area for every dorsal striatum region (ROI) analyzed. The size of each data point represents the mean Iba1+ cell area for every ROI. **F.** Bar graphs showing the mean Iba1+ cell count normalized to dorsal striatum area. 4-5 sections/rat, matched on Bregma coordinates, were used for quantification. Group mean ± 95% CI with individual values are shown. Statistical analysis by two-way ANOVA revealed a hemisphere effect (p < 001). Post-hoc test was performed with Tukey’s HSD. **p < 0.01, ***p < 0.001. Group numbers; saline n = 7 and XPro1595 n = 10. **Abbreviations.** Iba1, ionized calcium-binding adapter molecule 1; PC, principal component; rAAV6, recombinant adeno-associated virus serotype 6; α-Syn, α-Synuclein; PFFs, preformed fibrils; AI, artificial intelligence; PCA, principle component analysis; CI, confidence interval; ANOVA, analysis of variance; HSD, honestly significant difference; ROI, region of interest.

Since semantic segmentation allows for more detailed analysis at a cellular level, we performed a PCA to identify which cell characteristics were most affected by rAAV6-α-Syn+PFFs. The two first PCs explained 52.5% and 30.5% of the variation, respectively. The PCA biplot with loadings showed that that Iba1+ cell area (“Area (µm^2^)”) contributed the most to PC1, while Iba1+ cell area divided by Iba1+ cell perimeter (“Area:Perimeter”) contributed the most to PC2 (Fig. 4B).

Two-way ANOVA revealed a simple main effect on Iba1+ cell area for striatal hemisphere (F1, 30 = 62.7, p < 0.001), but not for treatment (F_1, 30_ = 1.73, p = 0.198), and no interaction effect (F_1, 30_ = 0.292, p = 0.593). Thus, rAAV6-α-Syn+PFFs caused a reduction in Iba1+ cell area in the ipsilateral vs contralateral striatum in both saline- (-83 µm^2^, 95% CI [-120, -42], p < 0.001) and XPro1595-(-72 µm^2^, 95% CI [-110, -38], p < 0.001) treated rats (Fig. 4C).

An increased Area:Perimeter ratio has been used as a characteristic of activated microglia (George et al., 2019; George et al., 2021; Stetzik et al., 2022). Two-way ANOVA showed an effect on Area:Perimeter for striatal hemisphere (F_1, 30_ = 6.91, p = 0.0134), but Tukey’s HSD post-hoc test did not show any statistically significant differences for ipsilateral vs contralateral striatum (Fig. 4D). There were no effects on Area:Perimeter for treatment (F_1, 30_ = 0.938, p = 0.341) or interaction between hemisphere and treatment (F_1, 30_ = 0.619, p = 0.438). However, rAAV6-α-Syn+PFFs caused an increase in striatal Iba1+ cell counts (hemisphere effect; F_1, 30_ = 39.0, p < 0.001), with similar effects in the ipsilateral vs contralateral hemispheres of saline- (66, 95% CI [21, 110], p = 0.0017) and XPro1595 (64, 95% CI [27, 100], p < 0.001) treated rats (Fig. 4E and F).

The same approach as for striatum was used to analyze Iba1+ cells in SN. PCA and PCA biplot for Iba1+ cells in SN showed similar results as those for striatum (Supplementary Fig. 2A and B). Two-way ANOVA showed an effect for hemisphere on Iba1+ cell area (F_1, 28_ = 9.14, p = 0.00531) and Iba1+ cell Area:Perimeter ratio (F_1, 28_ = 6.16, p = 0.0194), but post-hoc tests did not identify any significant differences between SN hemispheres (Supplementary Fig. 2C and D). There were also similar Iba1+ cell counts in ipsilateral and contralateral SN (Supplementary Fig. 2E and F). As for the striatum, no effect of XPro1595 treatment was seen for microglial phenotypes in the SN.

Indicative results from unilateral injection of rAAV-(-)+DPBS showed minor effects on Iba1+ cell area and count, but not Area:Perimeter, in the dorsal striatum (Supplementary Fig. 1G) and in SN (Supplementary Fig. 1H).

Analysis of MHCII immunoreactivity following rAAV6-α-Syn+PFFs injections showed a distinct increase of MHCII+ area in ipsilateral versus contralateral hemispheres in both saline and XPro1595 treated rats (Fig. 5A and B). Quantification of the normalized MHCII+ area showed a significant hemisphere effect in the dorsal striatum (F_1, 32_ = 159, p < 0.001) and in SN (F_1, 32_ = 53.8, p < 0.001) but no simple main effect of treatment or interaction effects. Post-hoc tests revealed that rAAV6-α-Syn+PFFs caused an increase in normalized percentage of MHCII immunoreactive area in both the dorsal striatum (saline (8.1%, 95% CI [5.9, 10], p < 0.001) and XPro1595 (6.0%, 95% CI [4.0, 8.0], p < 0.001), Fig. 5C) and in SN (saline (2.4%, 95% CI [0.63, 4.1], p = 0.00452) and XPro1595 (3.8%, 95% CI [2.2, 5.4], p < 0.001), Fig. 5D). A minor increase in MHCII+ immunoreactivity was observed in the dorsal striatum, but not the SN, following unilateral rAAV6-(-)+DPBS injection (Supplementary Fig. 1I).

**Figure 5.**
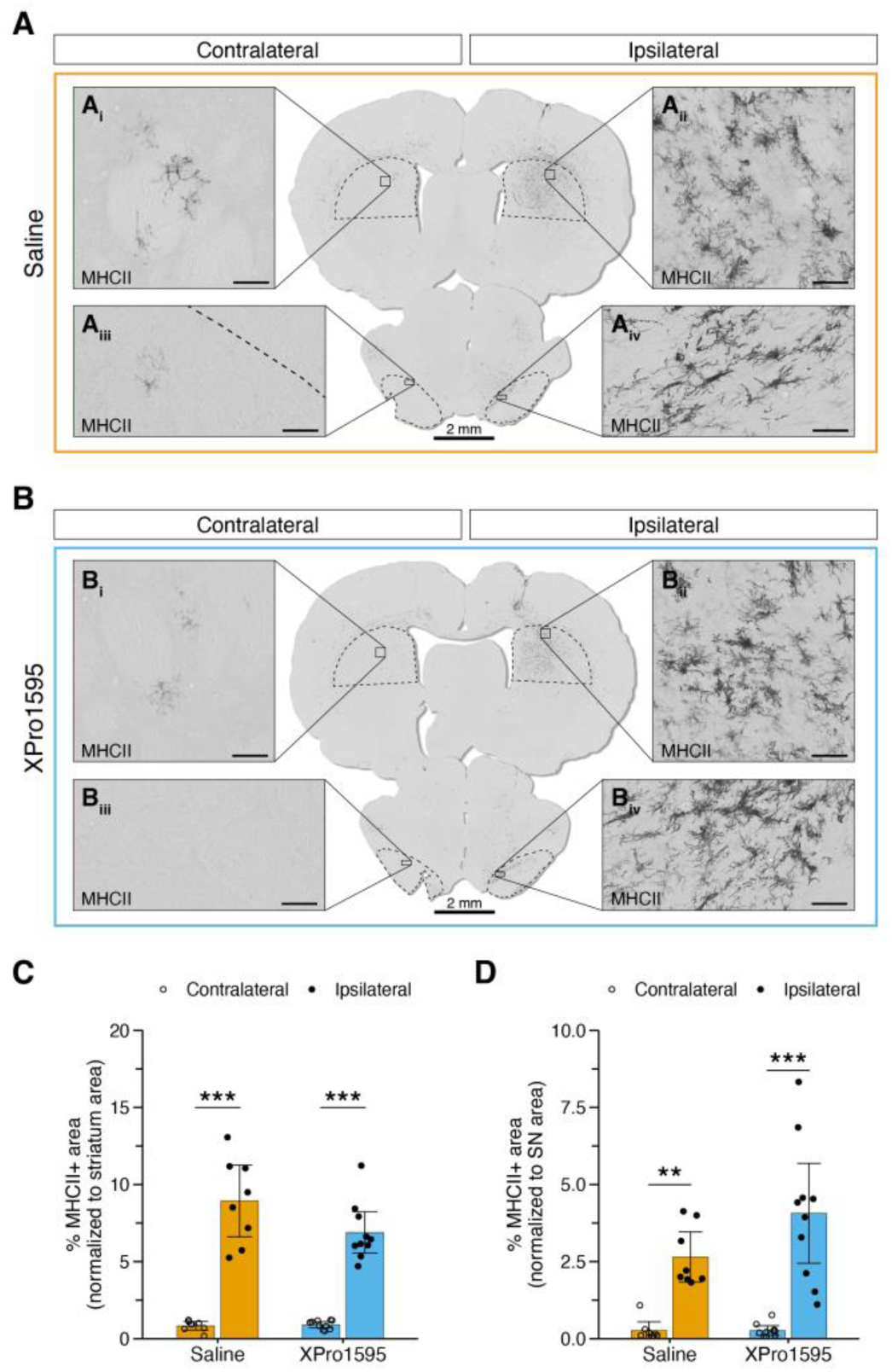
Systemic XPro1595 treatment does not reduce MHCII upregulation in dorsal striatum or SN following rAAV6-α-Syn+PFFs. A-B. Representative images of MHCII immunoreactivity in contralateral dorsal striatum (**A_i_** and **B_i_**) and SN (**A_iii_** and **B_iii_**) or in ipsilateral dorsal striatum (**A_ii_** and **B_ii_**) and SN (**A_iv_** and **B_iv_**) in saline treated control (**A**) or XPro1595 treated (**B**) rats. Representative ROI are indicated by dashed line. Scale bar (overview) = 2 mm. Scale bar (zoom) = 40 µm. **C.** Quantification of the percentage of MHCII+ area normalized to dorsal striatum area. 5 sections/rat, matched on Bregma coordinates, were used to calculate individual means. Group mean ± 95% CI with individual values are shown. Two-way ANOVA, showed an effect of hemisphere (p < 0.001). Post-hoc comparisons were made using Tukey’s HSD test. ***p < 0.001. **D.** Percentage of MHCII+ area normalized to SN area. 3-5 sections/rat were used to calculate individual means. Group mean ± 95% CI with individual values are shown. Two-way ANOVA, showed an effect of hemisphere (p < 0.001). Post-hoc comparisons were made using Tukey’s HSD test. **p < 0.01, ***p < 0.001. Group numbers; saline n = 8 and XPro1595 n = 10. **Abbreviations.** MHCII, major histocompatability complex class II; SN, substantia nigra; rAAV6, recombinant adeno-associated virus serotype 6; α-Syn, α-Synuclein; PFFs, preformed fibrils; ROI, region of interest; CI, confidence interval; ANOVA, analysis of variance; HSD, honestly significant difference.

To investigate microglia activation further we analyzed CD68, a marker for phagocytic activity. In the dorsal striatum, CD68 immunoreactivity was observed in the hemisphere ipsilateral to rAAV6-α-Syn+PFFs injections in both saline and XPro1595 treated rats, and to a lesser extent in the contralateral hemisphere (Fig. 6A). There was a hemisphere effect in the dorsal striatum (F_1, 32_ = 30.3, p < 0.001), with a similar increase in the ipsi-compared to contralateral hemisphere in saline (0.22%, 95% CI [0.066, 0.38], p = 0.00301) and XPro1595 (0.21%, 95% CI [0.066, 0.35], p = 0.00209) treated rats. (Fig. 6B).

**Figure 6.**
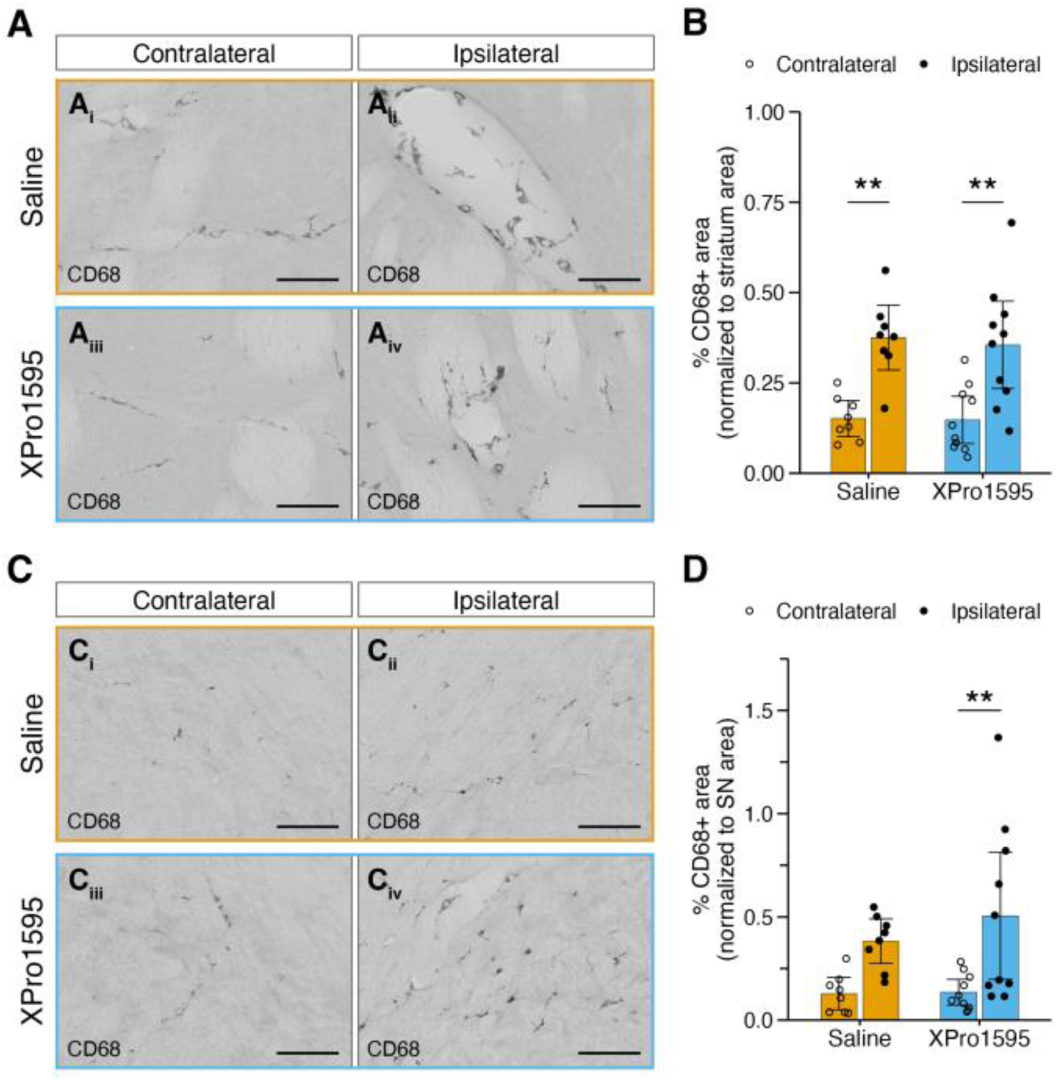
CD68 immunoreactive area is unaffected by sequestering of sTNF by systemic XPro1595 treatment following rAAV6-α-Syn+PFFs injections. **A.** Representative images of CD68+ signal in the dorsal striatum of saline and XPro1595 treated rats. Scale bar = 60 µm. **B.** Quantification of CD68+ area normalized to dorsal striatum area in saline and XPro1595 treated groups. 5 sections/rat, matched on Bregma coordinates, were used to calculate individual means. Group mean ± 95% CI with individual values are shown. Statistical analysis was performed using two-way ANOVA, showing an effect of hemisphere (p < 0.001). Post-hoc comparisons were made using Tukey’s HSD test. **p < 0.001. **C.** Representative images of CD68+ signal in the SN of saline and XPro1595 treated rats. Scale bar = 60 µm. **D.** Quantification of CD68+ area normalized to SN area, in saline and XPro1595 treatment groups. 3-5 sections/rat were used to calculate individual means. Group mean ± 95% CI with individual values are shown. Statistical analysis was performed using two-way ANOVA, showing an effect of hemisphere (p < 0.001). Post-hoc comparisons were made using Tukey’s HSD test. **p < 0.01. Group numbers; saline n = 8 and XPro1595 n = 10. **Abbreviations.** CD, cluster of differentiation; SN, substantia nigra; sTNF, soluble tumor necrosis factor; rAAV6, recombinant adeno-associated virus serotype 6; α-Syn, α-Synuclein; PFFs, preformed fibrils; CI, confidence interval; ANOVA, analysis of variance; HSD, honestly significant difference.

CD68 immunoreactivity was observed in both the contra- and ipsilateral SN (Fig. 6C), with a significant hemisphere effect (F1, 32 = 15.3, p < 0.001). Post-hoc analysis confirmed a statistically significant increase in the ipsi-compared to contralateral hemisphere in XPro1595 (0.37%, 95% CI [0.074, 0.67], p = 0.00982), but not in saline treated rats (Fig. 6D). XPro1595 treatment did not reduce the overall levels of CD68 immunoreactivity compared to saline. The results following rAAV6-(-)+DPBS injection indicated elevated levels of CD68 immunoreactivity in the ipsilateral dorsal striatum whereas CD68 levels remained low in the ipsilateral SN compared to contralateral hemispheres (Supplementary Fig. 1J).

### Astrocyte response was localized to the dorsal striatum following unilateral rAAV6-α-Syn+PFFs injection

Activated astrocytes, characterized by hypertrophy and upregulation of markers including GFAP, are seen in the SNpc of PD post-mortem brains (Ishida et al., 2006). To assess the astrocyte response to unilateral injection of rAAV6-α-Syn+PFFs, and the effect of sTNF inhibition using XPro1595, we investigated the GFAP+ area in the dorsal striatum and SN. In response to unilateral injection of rAAV6-α-Syn+PFFs, GFAP was upregulated in the ipsilateral dorsal striatum of both treatment groups (Fig. 7A). Two-way ANOVA confirmed a hemisphere effect (F_1, 32_ = 16.5, p < 0.001) but no effect from treatment (F_1, 32_ = 3.14, p = 0.086) or interaction (F_1, 32_ = 0.013, p = 0.91). The increased percentage of GFAP+ area in the ipsilateral vs contralateral striatum was similar for saline- (3.7%, 95% CI [0.11, 7.3], p=0.042) and XPro1595-treated (3.5%, 95% CI [0.28, 6.7], p = 0.029) rats (Fig. 7B).

**Figure 7.**
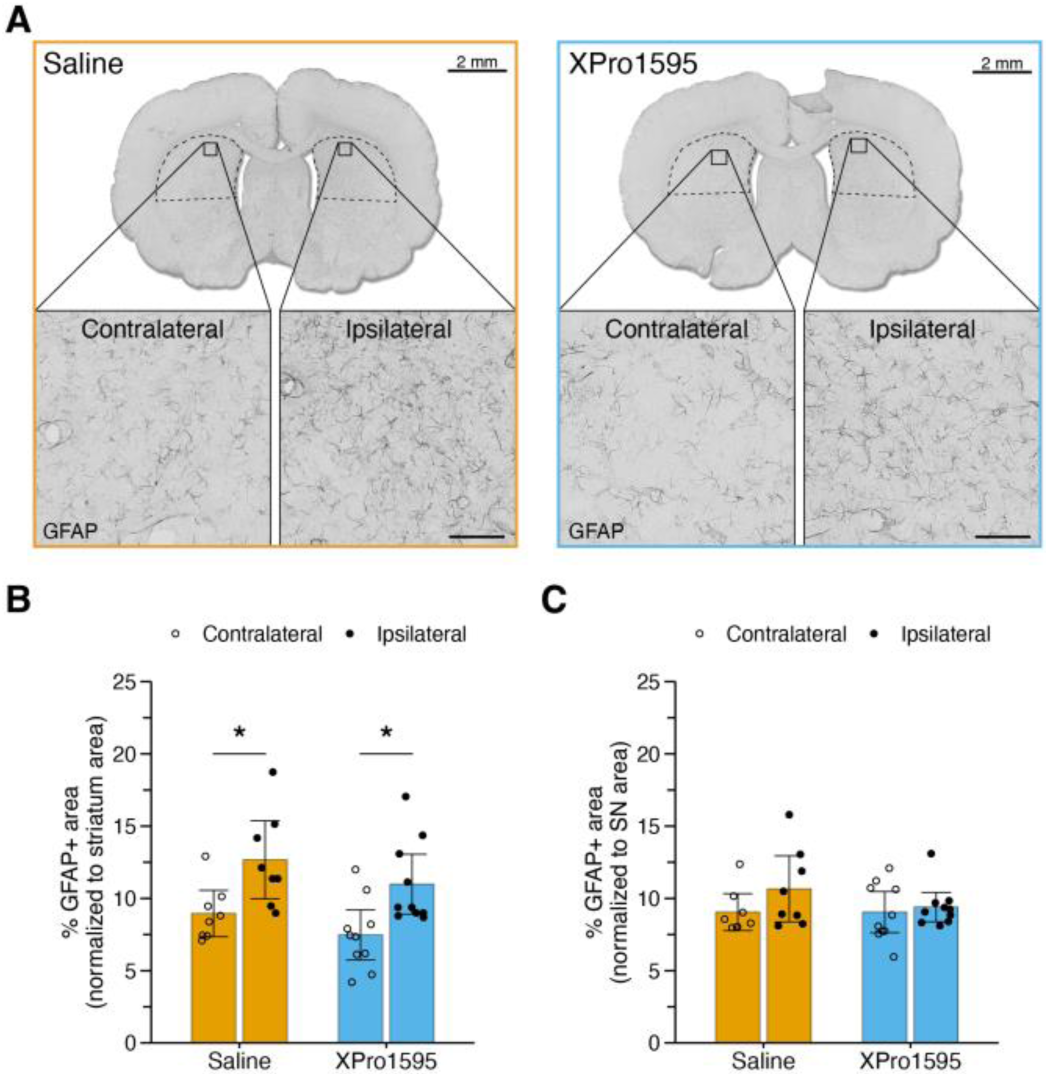
Unilateral rAAV6-α-Syn+PFFs injection caused astrocyte activation in dorsal striatum only. **A.** Representative images of GFAP+ signal in dorsal striatum of saline (left) and XPro1595 (right) treated rats following unilateral rAAV6-α-Syn+PFFs injection. Representative dorsal striatum region used for analysis is indicated by dashed outline. Scale bar (overview, top) = 2 mm. Scale bar (zoom, bottom) = 80 µm. **B.** Quantification of the percentage of GFAP+ area normalized to dorsal striatum area. 5 sections/rat, matched on Bregma coordinates, were used to calculate individual means. Group mean ± 95% CI with individual values are shown. Statistical analysis was performed using two-way ANOVA, showing an effect of hemisphere (p < 0.001). Post-hoc comparisons were made using Tukey’s HSD test. *p < 0.05. **C.** Bar graph showing mean percent GFAP+ area (normalized to SN area). 3-6 sections/rat were used to calculate individual means. Group mean ± 95% CI with individual values are shown. Statistical analysis was performed using two-way ANOVA. Post-hoc comparisons were made using Tukey’s HSD test. Group numbers; saline n = 8 and XPro1595 n = 10. **Abbreviations.** GFAP, glial fibrillary acidic protein; SN, substantia nigra; rAAV6, recombinant adeno-associated virus serotype 6; α-Syn, α-Synuclein; PFFs, preformed fibrils; CI, confidence interval; ANOVA, analysis of variance; HSD, honestly significant difference.

In contrast, rAAV6-α-Syn+PFFs had no effect on astrocyte activation, determined by GFAP signal in the SN (F_1, 32_ = 1.94, p = 0.174) (Fig. 7C), and no effect of treatment (F_1, 32_ = 0.918, p = 0.345) or interaction effect (F1, 32 = 0.932, p = 0.341) was detected. Following unilateral rAAV6-(-)+DPBS injection there was an indication of an increased percentage of GFAP+ area in the ipsilateral dorsal striatum and in SN (Supplementary Fig. 1K).

### XPro1595 treatment caused increased levels of proinflammatory cytokines in CSF but not in serum

To assess if sequestering native sTNF with XPro1595 impacted the levels of circulating cytokines in serum and CSF we analyzed IFNγ, IL-10, IL-13, IL-1β, IL-4, IL-5, IL-6, and CXCL1 by multiplexed ELISA.

In serum, only CXCL1 levels were detected (Supplementary Table 1). Serum CXCL1 levels were unaffected by systemic XPro1595 treatment. In CSF, 5/8 cytokines were detected (Supplementary Table 2). CSF levels of CXCL1, IFNγ and IL-10 remained unchanged, while IL-6 and IL-1β were elevated in the XPro1595-treated compared to the saline group (0.738 pg/ml, 95% CI [0.121, 1.36], p = 0.0233 and 1.16 pg/ml, 95% CI [0.133, 2.22], p = 0.0330, respectively).

Collectively these results indicate that sequestering sTNF with XPro1595 treatment caused minor, but detectable, increased levels of IL-6 and IL-1β in CSF whereas serum cytokine levels were low and unchanged following rAAV6-α-Syn+PFFs injection.

## Discussion

In this study, we investigated the effects of selective inhibition of sTNF on α-Syn-induced Parkinson-like pathology *in vivo* and found no treatment effects on neuroprotection, microglia or astrocytes. In previous studies, we found that congenic DA.VRA4 rats have elevated levels of sTNF in serum compared to the background strain DA, and that this correlates with increased neurodegeneration, pathological α-Syn spread and neuroinflammation (Fredlund et al., 2024; Jimenez-Ferrer et al., 2021). Unlike the temporal upregulation of the cellular immune response, sTNF levels were consistently higher in DA.VRA4 vs DA rats before, 4- and 8-weeks post rAAV6-α-Syn+PFFs, and were hypothesized to increase susceptibility to PD-like pathology (Fredlund et al., 2024).

The use of anti-TNF therapy to treat irritable bowel disease has been reported to lower the incidence of PD by 78%, suggesting anti-TNF treatment as a viable treatment strategy for PD (Peter et al., 2018). However, targeting TNF in neurodegenerative diseases has also been critiqued with the rationale that innate immunity in CNS is not well understood, and that TNF inhibition will not necessarily work in CNS even if it has been successful in adaptive immune disorders (Ransohoff, 2016). This rationale is supported by a recent Mendelian randomization study which reported that long-term use of TNF therapies blocking TNFR1 signaling would not have a protective effect on PD, whilst confirming a protective effect of TNFR1 blocking on Crohn’s disease and ulcerative colitis and an increased risk of multiple sclerosis (MS) (Kang et al., 2021). Although there are case studies were individuals taking TNF inhibitors have developed inflammatory neurological syndromes, including demyelination, it is still unknown if the effect is causal or pertains to genetic susceptibility (Gelfand, 2014; Solomon et al., 2011). Additionally, a phase II randomized placebo-controlled study using the TNF inhibitor lenercept, was halted due to exacerbated, rather than reduced, MS severity (1999). These adverse effects are likely due to the lack of specificity of current anti-TNF therapies targeting both sTNF/TNFR1 and tmTNF/TNFR2 signaling. The use of DN-TNF efficiently circumvents this issue.

Before the current study, there have been contradictory reports on whether TNF is associated with α-Syn pathology and neurodegeneration in α-Syn-induced PD models. The serum levels of sTNF and other proinflammatory cytokines were found to be unaffected after striatal injection of PFFs to mice, despite several changes in immune cell profiles preceding TH+ cell loss (Earls et al., 2019). On the other hand, elevated TNF levels were found in mouse striatum 8 weeks post PFFs injection (Karampetsou et al., 2017) and in mouse SNpc 4 weeks post rAAV2-mediated α-Syn overexpression (Theodore et al., 2008). Yet another study found higher TNF levels in rat striatum, but not SN, 8 weeks post rAAV2-mediated α-Syn overexpression in the SN (Chung et al., 2009). Of note, the studies reporting increased TNF in brain tissue have used ELISA- or qPCR-based quantification which does not discriminate between sTNF and tmTNF.

Previous reports from 6-OHDA-induced PD-like dopaminergic neurodegeneration in rats have clearly demonstrated a reduction in astrocyte activation, determined by GFAP intensity, after systemic XPro1595 treatment (Barnum et al., 2014). Here, we show that the astrocyte response is similar in XPro1595 treated rats compared to controls. However, it is important to consider the rat strains used; naïve DA.VRA4 rats are more inflammatory-prone compared to the DA background strain (Fredlund et al., 2024). Therefore, the trend observed in a reduced astrocyte response in both contra- and ipsilateral hemispheres in the dorsal striatum after XPro1595 treatment in the current study suggests that sTNF inhibition has a potential effect on astrocyte activity, in which case is inherent to the dorsal striatum only. We have previously shown that the DA.VRA4 rats have regional differences in inflammatory response, with more MHCII+ microglia in dorsal striatum specifically, but not in SN, compared to DA rats following unilateral rAAV6-α-Syn+PFFs injection (Jimenez-Ferrer et al., 2021).

Systemic sTNF inhibition using DN-TNF variants have demonstrated a significant reduction of Iba1+ cell numbers in the SNpc following 6-OHDA injection (Barnum et al., 2014; Harms et al., 2011), which was not observed in the current study. The microglia-mediated response was, similar to the astrocyte response, localized to the dorsal striatum and not the SN, and was not affected by the systemic XPro1595 treatment. These findings are supported by our previous studies showing more extensive microglia activation in the striatum rather than the SN in DA.VRA4 rats following rAAV-mediated α-Syn overexpression (Jimenez-Ferrer et al., 2017) or combined injections of rAAV6-α-Syn+PFFs (Jimenez-Ferrer et al., 2021). In the current study we utilized a deep CNN algorithm to analyze Iba1+ cells that allows for higher-throughput analyses of Iba1+ cells, making it possible to analyze all cells in the region of interest (Stetzik et al., 2022). In the current study the Iba1+ cell area was the microglial phenotype that changed the most in response to rAAV6-α-Syn+PFFs. It is important to highlight that using Iba1+ cell area as the main readout could contribute to false negative results if the microglia morphological changes were mainly attributed to changing from a resting to a hyper-ramified state, where the cell size would increase rather than decrease (Sanchez-Guajardo et al., 2010). In fact, following rAAV6-α-Syn injection, we have previously shown that the microglia morphology in the SN in DA.VRA4 rats is restricted to a shift from a resting to a hyper-ramified state (Jimenez-Ferrer et al., 2017). Therefore, it is possible that the microglia activation based on morphology of Iba1+ cells in the current study is underestimating the microglia response observed in the SN. Additional analyses of the inflammatory markers CD68 and MHCII revealed a clear neuroinflammatory response to exogenous α-Syn, with no signs of effects from XPro1595 treatment. Our findings are thus in contrast to previous research demonstrating that XPro1595 treatment can reduce the levels of both CD68 and MHCII during inflammatory responses. Direct XPro1595 administration to the site of injury has been shown to reduce CD68 mRNA levels during the peak response to spinal cord injury in mice (Lund et al., 2023). In a 5xFAD mouse model of Alzheimer’s disease the percentage of MHCII+ microglia and infiltrating macrophages were reduced following systemic XPro1595 treatment, as determined by flow cytometry (MacPherson et al., 2017). Additionally, in a MPTP monkey model of PD systemic XPro1595 treatment reduced CD68+ area in the lower colon and the total MHCII+ area in SN compared to vehicle-treated controls (Joers et al., 2020). Using the same model and experimental set up as in the current study, we have previously shown that the neuroinflammatory response is highest at four weeks after rAAV-α-Syn injection into SN (Fredlund et al., 2024). To fully elucidate a potential neuroinflammatory mitigation of the XPro1595 treatment, an earlier time point than the eight weeks used in the current study needs to be investigated. Collectively, the lack of a reduced neuroinflammatory response following XPro1595 treatment suggest that sTNF may not be the only driver of neuroinflammation following rAAV6-α-Syn+PFFs and that elevated sTNF levels are not associated to increased susceptibility of exogenous α-Syn-induced PD-like pathology in the DA.VRA4 rat strain.

In the current study the cytokine levels in serum and CSF following rAAV6-α-Syn+PFFs injection were generally low, which is consistent with what we have previously reported (Fredlund et al., 2024). Systemic XPro1595 treatment did not reduce any of the detectable cytokines analyzed in the current study. Instead, we observed a small increase in IL-6 and IL-1β levels in CSF samples following XPro1595 treatment. A previous study reported no change in IL-1β, but a reduction of CCL2 and TGFβ following XPro1595 treatment in the 5xFAD mouse model of Alzheimer’s disease (MacPherson et al., 2017). Although both IL-6 and IL-1β can be produced by activated microglia and astrocytes, the cellular source is not determined in the current study. Additionally, IL-6 and IL-1β may activate microglia and astrocytes (reviewed in (Erta et al., 2012; Shaftel et al., 2008)), thus, it is possible that the elevated IL-6 and IL-1β CSF levels observed in the current study counteracts a possible reduction in neuroinflammation following neutralization of native sTNF.

Although there was no statistical difference in TH+ axonal swellings between saline- and XPro1595 treated rats, the intra-group variation in the XPro1595 treated group was attributed to three rats, where the rat with the highest count accounted for almost one third of all TH+ axonal swellings in the group (data not shown). The reason behind this observation is unclear, but it is worth noting that the same rat had, by far, the most pS129+ α-Syn pathology in the dorsal striatum, but little neurodegeneration. Although this could be due to biological or methodological variation, it is also possible that despite extensive α-Syn- and TH+ axonal swelling pathology, the XPro1595 treatment postponed neurodegeneration by sequestering sTNF. The results can, however, not support this effect on a group level.

In the current study, we cannot rule out that XPro1595 treatment was initiated outside the potential therapeutic window. Previous studies using DN-TNF have shown that direct delivery into the SNpc by infusion was protective against dopaminergic neurodegeneration induced by both 6-OHDA and low-dose LPS (McCoy et al., 2006). Additionally, lentiviral expression DN-TNF improved dopaminergic neuronal survival upon codelivery with 6-OHDA (McCoy et al., 2008). Even a two-week delayed lentivirus delivery into the SNpc completely stopped the progressive dopaminergic neurodegeneration induced by 6-OHDA (Harms et al., 2011). In contrast, a two-week delay of initiating systemic treatment with XPro1595 following 6-OHDA was ineffective and highlights the importance of the route of administration and therapeutic window (Barnum et al., 2014). Even if rAAV-mediated transgene delivery causes an immediate immune reaction (Martino and Markusic, 2020; Muhuri et al., 2021), rAAV-α-Syn-induced neurodegeneration is relatively slow-progressing (Cenci and Bjorklund, 2020). In the current study we use a combination of rAAV6-α-Syn and PFFs, enabling a reduced rAAV6-α- Syn titer, since the PFFs delivery efficiently, rapidly, and robustly initiates PD-like pathology (Bjorklund et al., 2022; Cenci and Bjorklund, 2020; Thakur et al., 2017). Additionally, we use human α-Syn overexpression, which is less potent in initiating pathological responses compared to murine-derived α-Syn in a murine model system (Luk et al., 2016). In the current study we hypothesized that, based on the available literature regarding timing of DN-TNF delivery and the characteristics of our model system, that initiating treatment one week post rAAV6-α-Syn SNpc delivery would be within the therapeutic window. Since we have not characterized the response to rAAV6-α-Syn one week post SNpc delivery, it is possible that there are interfering pathological processes occurring already at one week, that renders the use of delayed sTNF inhibition ineffective. Alternatively, the addition of exogenous PFFs overcomes any protective effect of α-Syn transgene overexpression alone. To fully elucidate the potential of sTNF inhibition in α-Syn-induced PD models, initiating XPro1595 treatment at the time of transgene delivery needs to be evaluated.

## Conclusion

This study shows that delayed systemically administered second-generation selective sTNF inhibitor XPro1595 successfully reached the CNS, but failed to exert neuroprotective effects. We found no effects from XPro1595 treatment on dopaminergic neurodegeneration, α-Syn pathology or glial cell activation at 8 weeks post unilateral injection of rAAV6-α-Syn followed by striatal administration of PFFs. The lack of a reduced neuroinflammatory response following XPro1595 treatment may be explained by increased levels of IL-6 and IL-1β in CSF. It is of high importance for future investigations of immunomodulatory therapies in PD, and anti-TNF therapy in particular, to conclude if sTNF inhibition is effective on toxin-induced dopaminergic neurodegeneration only, and if certain aspects make robust α-Syn-induced PD models unresponsive to delayed sTNF inhibition.

## Author contributions

**Filip Fredlund**: Conceptualization, data curation, formal analysis, funding acquisition, investigation, methodology, project administration, validation, visualization, writing – original draft, writing – review and editing. **Claes Fryklund**: Data curation, investigation, methodology, validation, writing – review and editing. **Olivia Trujeque-Ramos**: Investigation, writing – review and editing. **Hannah A. Staley**: Data curation, investigation, validation, writing – review and editing. **Joaquin Pardo**: Methodology and validation. **Kelvin C. Luk**: Resources, writing – review and editing. **Malú G. Tansey**: Conceptualization, supervision, resources, writing – review and editing. **Maria Swanberg**: Conceptualization, funding acquisition, resources, supervision, writing – original draft, writing – review and editing.

## Acknowledgements

We want to acknowledge Catarina Blennow for assistance with IHC, the MultiPark AAV Vector Lab platform and Jenny G. Johansson for production of the rAAV constructs, Lund Stem Cell Center Imaging Facility and Emanuela Monni for access to the Olympus VS-120 virtual slide microscope, and INmune Bio Inc. for providing XPro1595.

## Data statement

The data supporting the findings of this study are available on request from the corresponding author.

