## Supplementary data for "Lack of neuroprotection after systemic administration of the soluble TNF inhibitor XPro1595 in an rAAV6-α-Syn+PFFs-induced rat model for Parkinson’s disease"

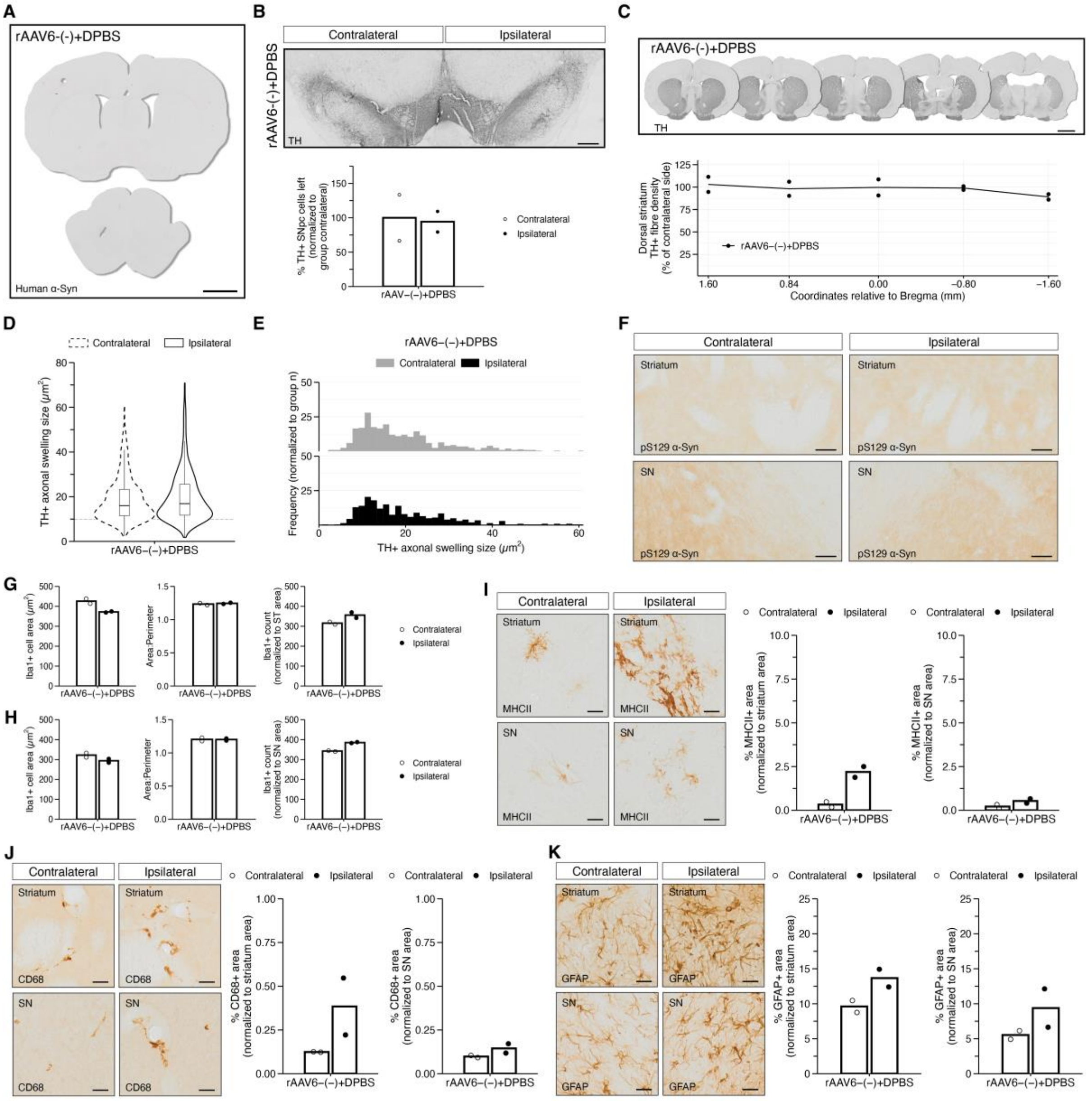


**Supplementary Figure 1. Indications that unilateral rAAV6-(-)+DPBS injection does not result in dopaminergic neurodegeneration, TH+ axonal swellings, α-Syn pathology, or marked glial cell activation. A.** Lack of human α-Syn expression eight weeks after unilateral rAAV-(-) injection in SNpc combined with striatal seeding with DPBS. Scale bar = 2 mm. **B.** Representative image of TH+ cells in ventral midbrain (top, scale bar = 400 μm) and quantification of the percentage of TH+ cells left in the SNpc normalized to the group contralateral (bottom). Bar graph showing group mean with individual values. **C.** Representative coronal sections stained for TH at various Bregma coordinates (top, scale bar = 2 mm). The TH+ fiber density was measured in the dorsal striatum. Quantification of the percentage TH+ fiber density remaining in the ipsilateral dorsal striatum after normalization to contralateral hemisphere, plotted against Bregma coordinates along the rostro-caudal axis (bottom). Group mean with individual values are shown. **D.** Violin plots of TH+ axonal swelling sizes (μm$^2$) based on the sum of TH+ axonal swellings in the dorsal striatum. **E.** Histogram plots of TH+ axonal swelling sizes (μm$^2$) for each hemisphere normalized to the group number. **F.** Representative images showing lack of pS129 α-Syn signal in contra- and ipsilateral striatum (top) and SN (bottom). Scale bar = 50 μm. **G-H.** Minimal microglia activation following rAAV6-(-)+DPBS injection in striatum (**G**) and SN (**H**) based on Iba1+ cell area (μm$^2$) and count. Bar graph showing group mean with individual values. **I.** Representative images of MHCII immunoreactivity in dorsal striatum and in SN (left, scale bar = 25 μm) and quantification of the percentage of MHCII+ area normalized to ROI area in dorsal striatum and in SN (right). Group mean and individual values are shown. **J.** Representative images of CD68 immunoreactivity in dorsal striatum and in SN (left, scale bar = 25 μm) and quantification of the percentage of CD68+ area normalized to ROI area in dorsal striatum and in SN (right). Group mean and individual values are shown. **K.** Representative images of GFAP immunoreactivity in dorsal striatum and in SN (left, scale bar = 25 μm) and quantification of the percentage of GFAP+ area normalized to ROI area in dorsal striatum and in SN (right). Group mean and individual values are shown. **A, B** and **G-K.** A total of 5 sections/rat, matched on Bregma coordinates, were used for quantification in dorsal striatum.

A total of 3-6 sections/rat were used for quantification in SN. **Abbreviations.** rAAV6, recombinant adeno-associated virus serotype 6; DPBS, Dulbecco's phosphate buffered saline; α-Syn, α-Synuclein; TH, tyrosine hydroxylase; SNpc, substantia nigra pars compacta; pS129, phosphorylated serine residue 129; SN, substantia nigra; Iba1, ionized calcium-binding adapter molecule 1; ST, striatum; MHCII, major histocompatability complex class II; CD, cluster of differentiation; GFAP, glial fibrillary acidic protein; ROI, region of interest.

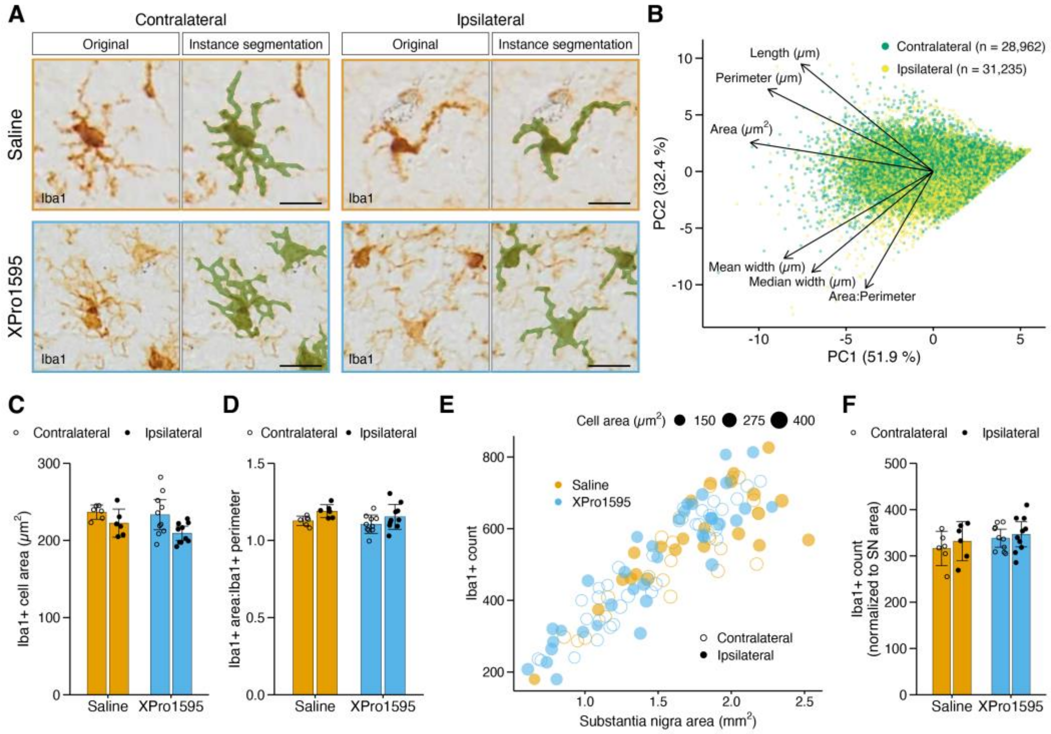


**Supplementary Figure 2. Microglia morphology in SN was unaffected by systemic XPro1595 treatmend following combined unilateral injection of rAAV6-α-Syn+PFFs. A.** Representative images of Iba1+ cells in the contralateral and ipsilateral SN in both groups. Instance segmentation overlaps shows how the AI model detected and outlined Iba1+ cells. Scale bar = 20 µm. **B.** PCA biplot with loadings for every instance segmentation event detected in SN, seperated by hemispheres. **C.** Bar graph showing the mean Iba1+ cell area across groups. 1-6 sections/rat were used to calculate individual means. Group mean ± 95% CI with individual values are shown. Statistical analysis was performed using two-way ANOVA, showing an effect of hemisphere ($p < 0.01$). Post-hoc comparisons were made using Tukey's HSD test. **D.** Quantification of the mean ratio of Iba1+ area divided by the Iba1+ perimeter across groups. 1-6 sections/rat were used to calculate individual means. Bar graph shows mean ± 95% CI with individual values. Two-way ANOVA showed a simple main effect of hemisphere ($p < 0.05$). Tukey's HSD was used for post-hoc comparison. **E.** Scatter plot showing the mean Iba1+ cell count against the area for every SN region (ROI) analyzed. The size of each data point represents the mean Iba1+ cell area for every ROI. **F.** Bar graphs showing the mean Iba1+ cell count normalized to SN area. 1-6 sections/rat were used for quantification. Group mean ± 95% CI with individual values are shown. Statistical analysis by two-way ANOVA revealed no simple main or interaction effects. Group numbers; saline n = 6 and XPro1595 n = 10. **Abbreviations.** Iba1, ionized calcium-binding adapter molecule 1; PC, principal component; SN, substantia nigra; rAAV6, recombinant adeno-associated virus serotype 6; α-Syn, α-Synuclein; PFFs, preformed fibrils; AI, artificial intelligence; PCA, principle component analysis; CI, confidence interval; ANOVA, analysis of variance; HSD, honestly significant difference; ROI, region of interest.

**Supplementary Table 1. Cytokine levels (pg/ml) in serum samples**

LLOQ and ULOQ (pg/ml) is specified in hard brackets after each specific cytokine; [LLOQ, ULOQ]. Data is presented as mean ± SD. Unpaired Welch's t-test was used to compare groups.

| Serum cytokines (pg/ml) | Saline (n = 8) | XPro1595 (n = 10) | Statistics |
|---|---|---|---|
| **IFNγ [3.85, 5290]** | ND | ND | - |
| **IL-10 [4.21, 25,900]** | ND | ND | - |
| **IL-13 [0.112, 1690]** | ND | ND | - |
| **IL-1β [0.407, 20400]** | ND | ND | - |
| **IL-4 [0.273, 1130]** | ND | ND | - |
| **IL-5 [2.19, 10200]** | ND | ND | - |
| **IL-6 [1.42, 13900]** | ND | ND | - |
| **CXCL1 [0.0103, 2680]** | 682 ± 390 | 558 ± 300 | ns |

**Abbreviations.** LLOQ, lower limit of quantification; ULOQ, upper limit of quantification; SD, standard deviation; IFNγ, interferon γ; ns, non-significant; IL, interleukin; ND, non-detected; CXCL1, C-X-C motif chemokine ligand 1.

**Supplementary Table 2. Cytokine levels (pg/ml) in CSF samples**

LLOQ and ULOQ (pg/ml) is specified in hard brackets after each specific cytokine; [LLOQ, ULOQ]. Data with parametric distribution is presented as mean ± SD. *Data with non-parametric distribution is presented as median (IQR). Unpaired Welch's t-test was used to compare groups with parametric data distribution. Statistically significant differences are presented as; difference in means, 95% CI [lower limit, upper limit], p-value. Mann-Whitney U test was used to compare groups with non-parametric data distribution.

| CSF cytokines (pg/ml) | Saline (n = 7) | XPro1595 (n = 7) | Statistics |
|---|---|---|---|
| **IFNγ [3.85, 5290]** | 2.81 ± 2.34 | 3.25 ± 2.31 | ns |
| **IL-10 [4.21, 25900]** | 1.50 ± 1.50 | 1.97 ± 0.850 | ns |
| **IL-13 [0.112, 1690]** | ND | ND | - |
| **IL-1β [0.407, 20400]** | 1.71 ± 1.02 | 2.88 ± 0.740 | 1.16 pg/ml, 95% CI [0.113, 2.22], p = 0.0330 |
| **IL-4 [0.273, 1130]** | ND | ND | - |
| **IL-5 [2.19, 10200]** | ND | ND | - |
| **IL-6 [1.42, 13900]** | 1.09 ± 0.600 | 1.83 ± 0.436 | 0.738 pg/ml, 95% CI [0.121, 1.36], p = 0.0233 |
| ***CXCL1 [0.0103, 2680]** | 26.2 (44.9) | 55.5 (35.3) | ns |

**Abbreviations.** CSF, cerebrospinal fluid; LLOQ, lower limit of quantification; ULOQ, upper limit of quantification; SD, standard deviation; IQR, interquartile range; CI, confidence interval; IFNγ, interferon γ; ns, non-significant; IL, interleukin; ND, non-detected; CXCL1, C-X-C motif chemokine ligand 1.
